# TEAD inhibition alters the lung immune microenvironment and attenuates metastasis

**DOI:** 10.1101/2025.02.26.639326

**Authors:** Raneen Rahhal, Marcel O. Schmidt, Maha Moussa, Tina Asemi, Rohith Battina, Amber Kiliti, Anton Wellstein, Anna T. Riegel, Ghada M Sharif

## Abstract

The Hippo pathway signaling mediated through YAP/TAZ, and the transcription factor TEAD is known to be involved in primary tumor progression. Here we report that novel TEAD inhibitors (iTEAD) cause a significant reduction in the outgrowth of lung metastases from triple negative breast cancer (TNBC) models mediated predominantly through changes in stromal immune signaling. TEAD inhibition did not affect the proliferation of TNBC cancer cells *in vitro* or the growth of the primary tumor *in vivo*. In normal mice that were treated with iTEAD in the absence of tumors, the lungs showed a decrease in pro-tumor inflammatory pathways. However, the IL12 signaling pathway was enhanced and its production from isolated lung tissue resident macrophages, but not bone marrow derived macrophages, was elevated. In syngeneic TNBC mouse models, inhibition of TEAD suppressed pro-tumor inflammation and the M2-like macrophage phenotype in lung tissues, and increased the infiltration of CD8+ T cells into the lung as well as Th1 CD4+ T cells, restoring an immune responsive microenvironment. iTEAD-treated T cells showed enhanced cytotoxicity and degranulation when co-cultured with cancer cells via increased IL-2 activity. Furthermore, TEAD inhibition or knockdown, enhanced T-cell macrophage crosstalk and anti-tumor activity in 3D tumorspheres which was reversed by IL12 neutralizing antibodies. Our data supports a multifaceted model of TEAD inhibition on the innate and adaptive immune cells as they respond to tumor cell signals and reveals an important stromal phenotype by which TEAD inhibitors could reverse immune suppression and eliminate seeded metastases in the lungs.

## Introduction

Metastatic triple negative breast cancer (TNBC) is an aggressive disease with short overall survival rate (1). Patients with metastatic TNBC are treated with a combination of chemotherapy, immunotherapy, as well as select targeted therapies, which only benefit a small subset of patients (2). During early stages of primary tumor development, cancer cells disseminate and seed secondary organs but remain undetectable for years after initial diagnosis (3). Metastatic disease accounts for over 90% of cancer-related mortalities (4). Therefore, it is imperative to develop strategies to prevent the outgrowth of micrometastases and inhibit metastatic progression.

During tumor progression, cancer cells use suppressive mechanisms to evade immune surveillance that ultimately inhibits immune cell anti-tumoral functions and promote tumor growth. The extent and composition of immune cell infiltration within a tumor can be indicative of patient treatment response (5). There have been many efforts towards reinvigorating the immune system, notably using anti-PD1/PD-L1 treatments to restore exhausted T-cell activity, and more recent attempts to induce macrophage polarization from an anti-inflammatory M2 state to a pro-inflammatory M1 state (6,7). While immunotherapies have shown varied success in patients with advanced disease, their efficacy in TNBC are limited due to the cold status of the immune microenvironment (2). A better understanding of the tumor microenvironment in TNBC is crucial for developing effective therapeutic strategies.

TEA domain (TEAD) transcription factors function during early development, tissue regeneration, and wound healing; however, they have also been implicated in tumorigenesis, metastasis, and drug resistance (8,9). TEADs primarily bind the well-established coactivator oncoproteins YAP and TAZ, which are downstream direct targets of the Hippo pathway, thus activating oncogenic transcription (10). We have previously showed that YAP/TEAD transcriptional program leads to progression of early-stage TNBC (11). Other studies have used YAP/TAZ overexpression mouse models to reveal their role as oncogenes and drivers of metastasis (12–14). Elevated expression of YAP and its downstream targets is correlated with a decreased survival in TNBC patients compared to hormone receptor positive and HER2+ patients (15). This data highlights an important role of YAP/TEAD regulated transcription in metastatic TNBC deeming it an attractive therapeutic target to halt metastasis and improve patient outcomes.

Preclinical studies have demonstrated that TEAD inhibitors can suppress tumor growth in YAP/TAZ-dependent cancers. Small molecules that inhibit TEAD autopalmitoylation and its binding to YAP show promising antitumor effects with multiple drugs currently in phase I clinical trials (VT3989-NCT04665206, ISM6331-NCT06566079, and IAG933-NCT04857372) (16–18). These inhibitors were originally tested on neurofibromatosis type 2 (NF2) negative or mutant cell lines, a tumor suppressor and upstream regulator of the Hippo pathway. These drugs showed anti-proliferative effects that were not seen on NF2 wildtype cell lines (19). Surprisingly, initial results from a clinical trial of VT3989 showed a benefit in patients with metastatic tumors regardless of their NF2 status (17) suggesting an alternative mechanism of action.

Evolving evidence suggest that YAP/TEAD may impact the immune stroma in solid tumors (20). In prostate cancer, activated YAP signaling increased CXCL5 expression which recruited myeloid-derived suppressor cells (MDSC), promoting immune evasion (21). YAP expression was associated with M2 tumor-associated macrophage polarization and disease progression in colon cancer (22). Additionally, the hippo pathway has been implicated in T cell development and function (23,24). YAP/TAZ transcription contributes to immunotherapy resistance as it can increase the expression of PD-L1 in melanoma, lung and breast cancers (25,26). Importantly, YAP expression is essential for tumor immune suppression by Treg cells and attenuates CD8 T cell immunity (27–29). PRKCI promoted immune suppression in ovarian carcinoma through downstream function of YAP that enhanced myeloid-derived suppressor cells and inhibited cytotoxic T cell infiltration (30). Collectively, this data demonstrates the potential to target downstream targets of the Hippo pathway to activate immune cells and hinder tumor progression.

Here we showed a novel anti-metastatic effect of TEAD inhibitors in TNBC mouse models through direct stromal effects irrespective of the NF-2 status of the primary tumor. We defined the function of TEAD-regulated transcription in the immune microenvironment of the lungs during metastatic progression and its impact on the outgrowth of metastases from primary mammary tumors. TEAD inhibition by a small molecule inhibitor or reduction by RNA interference reversed pro-tumor activation of macrophages, reduced inflammatory pathways and enhanced CD8 T cell cytotoxicity.

## Results

### TEAD inhibition reduces lung metastases but does not affect the primary tumor growth

To determine the effect of TEAD inhibition on triple-negative breast cancer (TNBC) progression, we utilized an early-stage breast cancer cell line MCFDCIS (DCIS) that metastasizes to the lungs when a small population of cells (DCISΔ4) expresses an isoform of the transcription coactivator amplified in breast cancer 1 (31). We injected half a million DCIS:DCISΔ4 cells into the mammary fat pad of NOD/SCID mice. Mice were treated daily with 30mg/kg of VT107, a pan TEAD inhibitor (32), or DMSO as a vehicle control, via oral gavage for 4 weeks. Tumors were surgically removed at week 4 and lungs were monitored for metastases for another 4 weeks without receiving additional treatment (Fig 1A). Mouse body weights were measured weekly throughout the course of treatment to monitor any adverse health events due to treatment (Fig S1A). Primary tumor size was similar between vehicle and VT107 treated mice (Fig 1B) which was consistent with *in vitro* data showing no significant change in DCIS proliferation in the presence of VT107 at varying concentrations (Fig S1B). All mice in the vehicle treated group developed lung metastases as previously reported (31). In contrast, the VT107-treated group showed no overt metastases in 4 out of 5 mice (Fig 1C). However, detection of human actin in lung tissues by qPCR, relative to mouse actin levels, indicated the presence of DCIS cells in the lungs of VT107-treated mice (Fig S1C). Immunofluorescent staining of the DCIS:DCISΔ4 cells confirmed the presence of single cancer cells in the lungs that failed to grow into overt metastases in the VT107-treated group (Fig 1D). This data suggests that TEAD inhibitors could directly influence the microenvironment to hinder the metastatic outgrowth of cancer cells that have already seeded the lungs. RNA was extracted from tumor and lung tissues to further examine host changes and determine gene expression and signaling pathways affected by VT107. Tumor tissues are predominantly comprised of human cancer cells; therefore, these RNA sequencing reads were aligned to the human transcriptome (Fig. S1D). RNA sequencing reads from the lungs were also aligned to the mouse transcriptome to shed light on the host tissue changes in response to treatment in the presence of seeded cancer cells (Fig S1E). Overall, the number of differentially expressed genes in the primary tumors were less than those observed in the lungs. Ingenuity pathway analysis of gene expression changes in the mouse lung stroma revealed an upregulation of myeloid cell activating pathways, acute phase response signaling and PD-L1 immunotherapy pathways. Signaling pathways implicated in pulmonary and hepatic fibrosis, cardiac hypertrophy and liver injury were downregulated (Fig 1E). These observations indicate a shift in the inflammatory phenotypes in the microenvironment which could be responsible for the halt in metastatic outgrowth of seeded cancer cells in the treated group. Given that many of the altered pathways pertain to immune functions, we next sought to investigate the impact of TEAD inhibition in an intact immune-competent setting.

**Figure 1:**
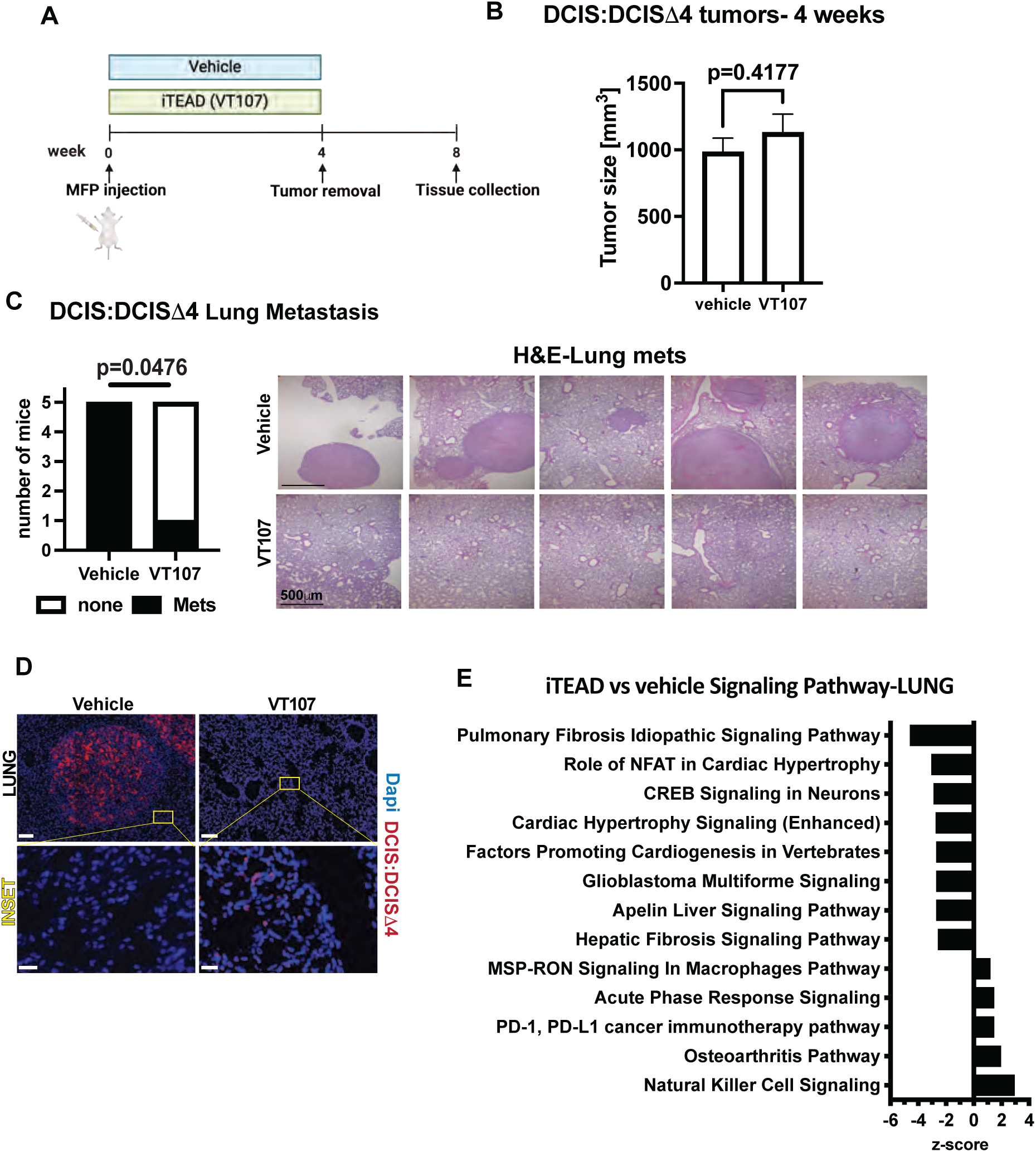
TEAD inhibition reduces lung metastases but has no effect on primary tumor growth. A) Experimental scheme of mammary fat pad (MFP) injection of DCIS:DCISΔ4 cells in NOD/SCID mice. Oral gavage with 30mg/kg of the iTEAD VT107 or DMSO vehicle started at the same time as MFP inoculation and terminated after primary tumor removal. B) Primary tumor volumes were measured after surgical resection at 4 weeks. (Mean±SD), student t-test, n=5 mice per group). C) Quantification of the number of NOD/SCID mice with lung metastases determined by histological analysis of lung tissue sections stained with hematoxylin and eosin (H&E) in VT107 treated and control mice. Representative fields of lung sections show metastatic lesions. (Fisher’s exact test). D) Immunofluorescent staining of DCIS:DCISΔ4 cell mix (red) in overt metastases or single cells in the lungs of mice that were treated with vehicle or VT107, respectively. The lower inset shows a magnified view of the lung tissue where single cancer cells were detected in VT107-treated mice. Scale bar is 100μM and 20μM in inset. E) Ingenuity pathway analysis (IPA) showing altered signaling pathways in the lungs of VT107-treated versus vehicle treated mice that have a p < 0.05 and a |z-score| > 1.3. The z-score indicate the directionality of the altered pathway, positive z-score value indicates upregulation while a negative value indicates downregulation of a signaling pathway. RNA-sequencing reads from whole lung tissues were aligned to the mouse transcriptome to analyze changes in the lung stroma only.

### TEAD inhibition alters the alveolar macrophage and monocyte populations in the lungs of naïve immunocompetent mice and activates IL12 signaling

To investigate how TEAD inhibition influences the immune microenvironment in normal lungs of immune-competent mice, we treated C57BL/6 mice with 30mg/kg of VT107 or vehicle control via oral gavage for 2 weeks. Lung, spleen and blood tissues were harvested for immune cell analysis and RNA-sequencing (Fig 2A). In the lungs, myeloid cell analysis by flow cytometry revealed a decrease in the overall population of monocytes, but an increase in tissue-resident alveolar macrophages (AM) (Fig 2B). No significant change in total macrophages, dendritic cells, eosinophils or neutrophils was observed in analyzed tissues (Fig S2A). Flow cytometry analysis of the lymphoid cell population in the lungs showed an increase in CD3+ cells, but not specifically in the CD4+ or CD8+ T cell populations (Fig S2B). These analyses suggest that VT107 treatment expanded the T cell population in the lungs, but in the absence of tumor cells, no specific clonal expansion occurred. RNA from the lungs and spleens were extracted for bulk RNA-sequencing. RNA-seq data was analyzed by Cibersortx (33) to deconvolute the abundance of cell types in the lung tissue using single-cell RNA-seq datasets. Consistent with our flow cytometry data, the monocyte fraction was reduced upon VT107 treatment and the tissue resident myeloid cells’ fraction that includes alveolar macrophages was expanded (Fig 2C). Gene expression analysis of the lungs showed a significantly higher number of downregulated genes compared to upregulated genes while there were minimal changes in the spleen (Fig S2C). Ingenuity pathway analysis of the lung gene expression data revealed overall reduction in inflammation with a decrease in cytokine storm, S100 family, wound healing and fibrosis signaling pathways, however, there was an increase in the IL12 signaling pathway (Fig 2D) which has been implicated in antitumor inflammation (34,35). Next, we sought to determine the specific effects of iTEAD treatment on lung resident versus bone-marrow derived macrophages as the dynamics of these cell populations are crucial for the establishment of a metastatic niche (36,37). We retrieved macrophages from the lungs by isolating F4/80+ cells, a specific marker for murine macrophages, by FACS (fluorescence-activated cell sorting) as well as bone marrow-derived macrophages from the femurs of naïve C57BL/6 mice that were then differentiated *in vitro* using macrophage colony stimulating factor (M-CSF). Macrophages were treated with 1μM of VT107 for 24 hours before the conditioned media was collected and analyzed on a cytokine array. Lung resident macrophages secreted a significantly higher level of IL12 and IL4 upon VT107 treatment, whereas iTEAD had no effect on IL12 and reduced IL4 levels in bone marrow-derived macrophages. There was no change in the level of the other cytokines detected (Fig. 2E and 2F). This data further illustrates the differential effect of TEAD inhibition on lung resident versus infiltrating monocyte/macrophage populations.

**Figure 2:**
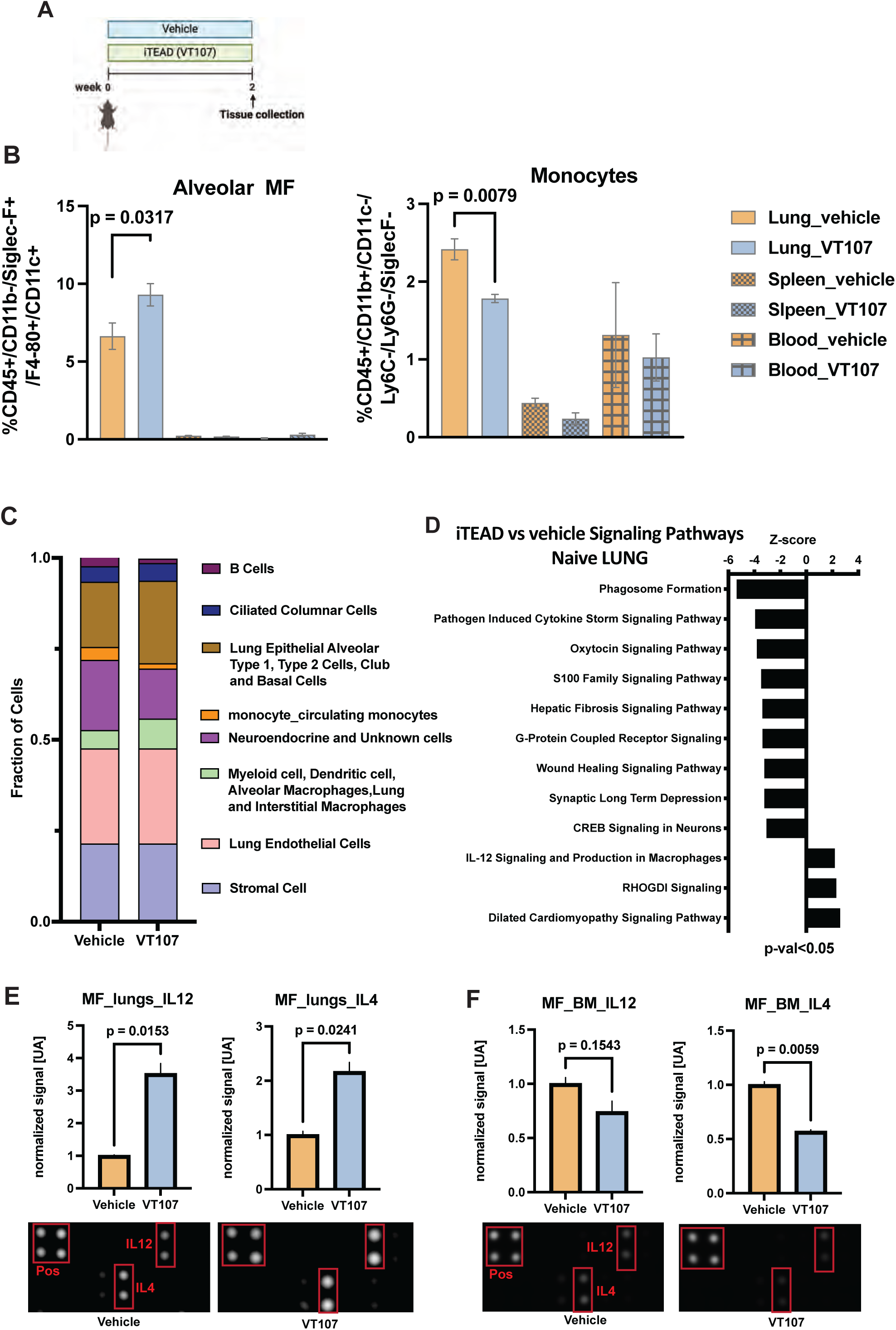
TEAD inhibition alters the alveolar macrophage and monocyte populations in the lungs of naïve immunocompetent mice and activates IL12 signaling. A) Experimental scheme of C57/Bl6 mice that were treated with 30mg/kg of the iTEAD VT107 or vehicle for two weeks before tissue collection. n=5 mice per group. B) Flow cytometry analysis of alveolar macrophages and monocytes in the lung, spleen and blood of naïve mice treated with VT107 or vehicle. (Mean±SEM, student t-test). C) CIBERSORTx (33) analysis of mRNA from lungs of VT107-treated and vehicle-treated groups showing changes in the abundance of tissue cell types. D) Ingenuity pathway analysis (IPA) showing differences in signaling pathways in the lungs of VT107-treated versus vehicle treated mice that have a p < 0.05 and a |z-score| > 2. E) Blot of cytokine array and quantified data showing changes in secreted factors in the conditioned media of isolated lung macrophages (MF_Lungs) treated with vehicle or 1μM VT107 for 24 hours. (Mean ± SEM, student t-test). F) Blot of cytokine array and quantified data showing changes in secreted factors in the conditioned media of isolated bone marrow macrophages (MF_BM) treated with vehicle or 1 μM VT107 for 24 hours. (Mean ± SEM, student t-test).

### TEAD inhibition repressed M2-like macrophage phenotype and enhanced their crosstalk with cancer cells

To further study the direct effects of TEAD inhibitors on macrophages, we utilized THP1, a human monocytic leukemia cell line that can differentiate into macrophages upon phorbol 12-myristate-13-acetate (PMA) treatment (38). iTEAD treatment on these macrophages showed an increase in IL12 and CD80 gene expression levels and a decreased expression of the known YAP/TEAD target genes CTGF, S100A8 and S100A9 (Fig S2D). To determine if knockdown of TEAD mimicked iTEAD effects, THP1 cells were transduced with shRNAs targeting TEAD 1, 3 and 4 or GFP as a control. Knockdown was confirmed by western blot with a pan TEAD antibody (Fig S2E). TEAD knockdown THP-1 macrophages migrated and attached to tumorspheres embedded in matrix compared to no effect in control macrophages (Fig. S2F and S2G). TEAD knockdown in THP1 macrophages decreased inflammatory cytokines CXCL2, 5, and 6 (Fig. 2H) and signaling pathways related to inflammatory diseases, such as rheumatoid arthritis (Fig.S2I), and inflammatory upstream regulators such as LPS and TNF (Fig.S2J). The top upregulated pathway in THP1 macrophages by TEAD knockdown was related to the immune interactions between lymphoid and non-lymphoid cells, a process necessary to mount an immune response (Fig.S2I) and is consistent with the enhanced crosstalk we observed between tumorspheres and shTEAD THP1 macrophages (Fig. S2F and S2G). The score for M2-like upstream regulators, OSM (39) and IL17A (40) was decreased by TEAD knockdown while an immune modulator used to enhance immunotherapy, NC410, was increased (41) (Fig S2J). Taken together, the response to iTEAD treatment or TEAD knockdown is a global anti-inflammatory phenotype in THP1 macrophages and repression of the M2-like macrophage phenotype, both of which oppose the establishment of a hospitable metastatic niche (42–44).

### TEAD inhibition suppresses pro-tumor inflammation, represses pro-tumor macrophages, and increases T cell populations in the lung of immunocompetent mice

The anti-metastatic effect of TEAD inhibition in NOD/SCID mice was related to immune signaling changes (Fig 1). iTEAD also impacted the immune microenvironment in the lungs of immunocompetent tumor naïve mice (Fig 2). Therefore, we next sought to determine the impact of iTEAD on tumor growth and lung metastases in an immune competent mouse model. We injected E0771 cells (45), into the mammary fat pad of C57BL/6 mice. When primary tumors became palpable at 3 weeks, mice were treated daily with 30mg/kg of VT107 or vehicle control via oral gavage. There was no impact on overall mice weights due to iTEAD treatment (Fig S3A). Primary tumors were surgically removed at week 4 and mice were euthanized at week 6 to collect tissues and inspect lungs for metastases (Fig 3A). Similar to our results in the DCIS:DCISΔ4 experiment in NOD/SCID mice (Fig 1), there was no difference in the primary tumor size between treatment groups (Fig 3B). E0771 cell proliferation *in vitro* was not affected by VT107 treatment (Fig S3B). Despite the early termination of the experiment due to recurrent mammary tumors (Fig S3C), histopathological assessment of lung sections from VT107-treated mice showed significantly less incidence of metastases compared to the vehicle treated group (Fig 3C). To investigate the tissue specific changes of iTEAD treatment, gene expression changes between treatment groups in lung and tumor tissues were analyzed by RNA-sequencing. A higher number of gene changes was observed in the lungs compared to the primary tumors (Fig. S3D). Pathway analysis showed a decrease in disease-related inflammation pathways such as hepatic and lung fibrosis and cardiac hypertrophy, and an activation of LXR/RXR anti-inflammatory pathway in the lung tissues (Fig. 3D). Consistent with our data in naïve mice (Fig 2), the resident alveolar macrophage population showed a small increase in the treatment group (Fig S3E) while the monocyte population followed a downward trend (Fig S3F). Importantly, CD86, a marker of M1-like macrophages, was significantly enriched in the macrophage population while CD206, a marker of M2-like macrophages, was significantly reduced (Fig. 3E and S3G). This shift was also significant in the alveolar macrophage population with iTEAD treatment (Fig. 3F). There was no difference in the CD68 positive cells in the tumors, a pan marker of monocytes and macrophages (Fig. S3H). Additionally, there was an increase in the number of CD8+ T cells infiltrating within the lungs, but not the primary tumors (Fig 3F and Fig S3H). Notably we observed CD8+ T cells in the metastatic lesions of the VT107-treated group whereas a minimal number of CD8+ T cells were observed in the metastases from the control group (Fig 3G). To determine early changes in the T cell population in a different syngeneic model *in vivo*, BALB/c mice were treated daily with 30mg/kg of VT107, or vehicle control via oral gavage for 15 days. On day 5, 4T1, a triple negative mouse mammary carcinoma cell line syngeneic to BALB/c mice (46), was injected into the tail vein to seed into the lungs (Fig. 3H). Lung and spleen tissues were processed and analyzed by flow cytometry 10 days post inoculation. There was no difference in the overall abundance of CD4+ and CD8+ T cell populations at this time point between treatment groups. However, the helper 1 T cell (Th1) population was increased in VT107-treated mice, but not other CD4+ T cell subtypes (Fig. S3I). A significant increase in T-bet staining, a marker of Th1 CD4+ cells(47), confirmed an enhanced Th1 cells’ presence in the lungs of iTEAD-treated mice (Fig. 3I). The alveolar macrophage population was increased in the lungs by iTEAD treatment consistent with what was observed in the tumor naïve mice (Fig 2) while the monocyte population decreased in the spleens (Fig. S3J). Lungs examined at day 15 (Fig 3H) had no detectable metastases in either treatment groups. However, another cohort of mice, terminated at day 21, showed a reduction in lung metastasis in response to iTEAD treatment (Fig. S3K and S3L). Thus, in both our TNBC syngeneic mouse models, iTEAD treatment reduced lung metastasis and shifted macrophages to an anti-tumor phenotype as well as increased T cell subpopulations in the lung. M1-like macrophages play a critical role triggering the proliferation and differentiation of T cell subsets that can eliminate tumor cells while M2-like macrophages suppress T cell proliferation (48–50). Thus, the interplay between macrophage and T cells after iTEAD treatment is likely to play a role in the increase in CD8+ T cell and Th1 CD4+ cell populations in the lungs.

**Figure 3:**
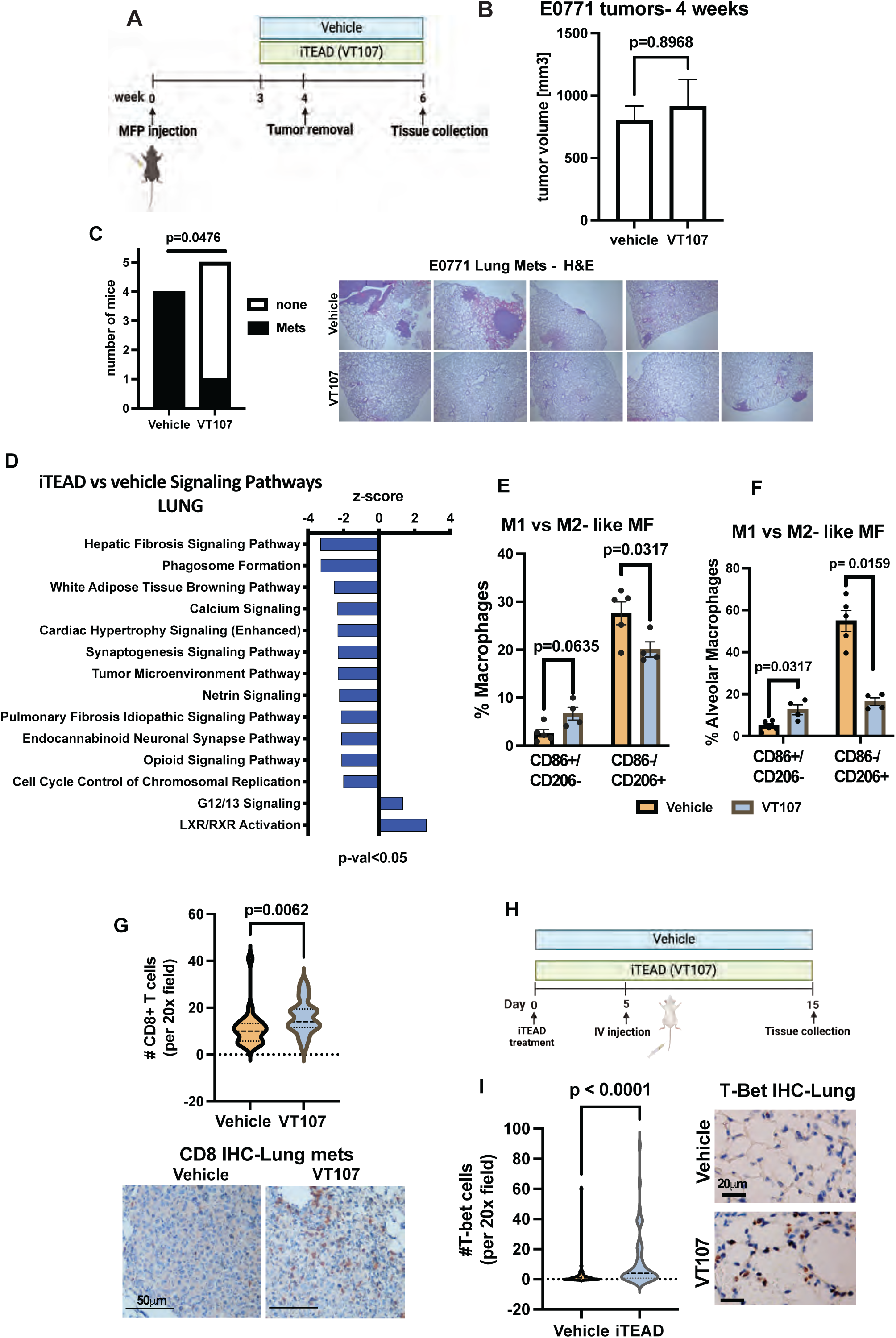
TEAD inhibition suppresses pro-tumor inflammation, activates anti-tumor macrophages and increases T cell populations in the lung of immunocompetent mice. A) Schematic of mammary fat pad (MFP) injection of E0771 cells in C57/Bl6 mice. Oral gavage with 30mg/kg of the iTEAD VT107 or vehicle started when tumors were palpable (3 weeks). B) Primary tumor volumes were measured after surgical removal at 4 weeks. (Mean±SD), student t-test, n=4-5 mice per group). C) Quantification of the number of C57/Bl6 mice that developed lung metastases determined by the histopathological analysis of lung tissue sections stained with H&E in the VT107-treated versus control group. Representative images of lung sections showing presence or absence of metastases. (Fisher’s exact test). D) IPA showing altered signaling pathways in the lungs of VT107-treated versus vehicle-treated group that have a p< 0.05 and a |z-score| > 1.3. E) Flow cytometry analysis showing percent of M1-like (CD86+) vs M2-like (CD206+) cells relative to the total macrophage population in the lungs of VT107 and vehicle-treated mice. (Mean±SEM, student t-test). F) Flow cytometry analysis showing percent of M1-like (CD86+) vs M2-like (CD206+) cells relative to the total alveolar macrophage population in the lungs of VT107 and vehicle-treated mice. (Mean±SEM, student t-test). G) IHC quantification for CD8 T cells in lung sections from vehicle vs iTEAD-treated mice. (Mean±SEM, Mann Whitney t-test, n=10 fields/lung). Representative images show differential infiltration of CD8+ T cells into lung metastases between treatment groups. H) Schematic of BALB/c mice that were injected intravenously (IV) with 4T1 cells and treated with either vehicle or 30mg/kg VT107 daily by oral gavage. I) IHC quantification of T-bet+ cells in the lungs of vehicle vs iTEAD-treated mice. (Mean±SEM, Mann Whitney t-test, n=10 fields/lung, n=5 mice per group). Representative images show the absence or presence of T-bet+ cells in lungs of vehicle or VT107 treated mice respectively.

### TEAD inhibition enhances antitumor T cell cytotoxicity and Th1 cell activation

To examine whether TEAD inhibition had a direct effect on T cells, we isolated T cells from the spleens of naïve mice, stimulated them overnight with anti-CD3 and anti-CD28 antibodies, followed by treatment with 1 μM of VT107 (Fig 4A). To test changes in their cytotoxic effect on cancer cells, 4T1 cells were plated on an array with an electrode-coated bottom to monitor changes to cell impedance, termed cell index, in real time. This measurement reflects cancer cell attachment as they proliferate or detachment as they die. After 4T1 cells attached, VT107-treated T cells were added to the cancer cells. There was a significant reduction in the cell index of 4T1 cells co-cultured with VT107-treated T cells compared to vehicle-treated T cells (Fig 4B). This effect was also observed on E0771 cancer cells (Fig S4A). In 3D tumorspheres, the number of 4T1 cells per sphere was significantly reduced when co-cultured for 72 hours with T cells that were pretreated with VT107 compared to vehicle (Fig. 4C). Cleaved caspase-3 immunohistochemistry (IHC) staining, a marker of apoptosis, was increased in 4T1 spheres with VT107-treated T cells (Fig. 4D). Additionally, VT107-treated T cells co-cultured with 4T1 cells in 2D for 72 hours showed a significant increase in CD107a, a marker of T cell degranulation which is indicative of cytotoxicity (51) (Fig. 4E). To identify changes in cytokine production by T-cells as a result of VT107 treatment, conditioned media was collected and assessed on a cytokine array. Levels of IL2, a cytokine produced by activated T cells (52), increased while IL13 and IL6 levels decreased (Fig S4B). Blocking of IL2 by a neutralizing antibody reversed the cytotoxic effect of VT107-treated T cells (Fig. 4F). This data demonstrates that TEAD inhibition has a direct effect on T cell activation and cytotoxicity via IL2 action.

**Figure 4:**
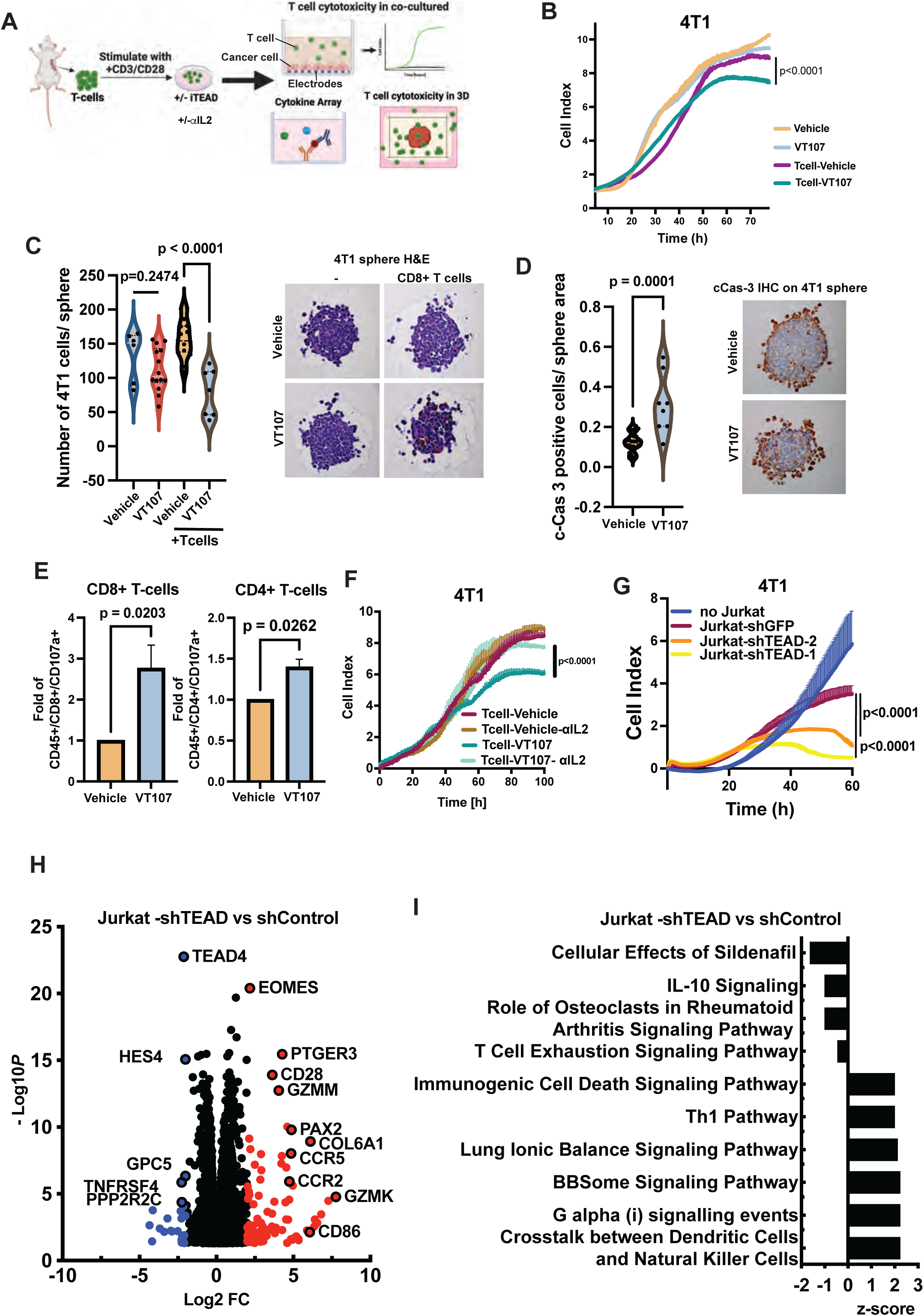
TEAD inhibition enhances T cell cytotoxicity and Th1 cell activation. A) Schematic of T cell isolation and downstream assays. T cells were isolated from naïve BALB/c mice spleens and pre-treated with 1μM VT107 or vehicle for 24hrs prior to assays. B) Real-time Cellular Analysis (RTCA) of 4T1 cell proliferation as measured by electric impedance sensing of attached cells (cell index) over time. T cells treated with vehicle or 1μM VT107 for 24 hours were added to cancer cells to monitor their cytotoxicity. (Mean±SD, student t-test). C) Quantification of 4T1 cell number in spheres after 72 hours co-culture with T cells that were pretreated with vehicle or 1μM VT107 for 24 hours. Treatment continued at time of co-culture. (Mean±SD, student t-test). Representative H&E images are shown. D) Quantification of cleaved caspase 3 (cCas-3) positive 4T1 cells normalized to average sphere size in each group. 4T1 spheres were co-cultured with T cells for 72 hours following 24-hour pretreatment with vehicle or 1μM VT107. (Mean±SD, student t-test). Representative IHC images are shown. E) Flow cytometry analysis showing change in CD107+ T cells, a degranulation marker. T cells isolated from the spleen were co-cultured with 4T1 cancer cells and treated with either vehicle or 1μM VT107 for 72hrs. F) RTCA of 4T1 cancer cells measured by electric impedance sensing of attached cells (cell index) over time. IL2 neutralizing antibody (50ng/ml) was added with the T cells as indicated. (Mean±SD, student t-test). G) RTCA of 4T1 cells was measured by electric impedance sensing of attached cells (cell index) over time. Jurkat cells that harbor shControl or shTEAD were added to the cancer cells to monitor their cytotoxicity. (Mean±SD, student t-test). H) Volcano plot of the altered gene expression in Jurkat cells that harbor shControl or shTEAD. Red and blue circles indicate regulated genes with −Log p-value >1.3 and |log_2_FC|=2. A full list of changed genes in supplementary table 2. I) IPA showing altered signaling pathways in shTEAD versus shControl Jurkat cells that have a p-val < 0.05.

A direct role of TEAD on T cell activation was also confirmed in Jurkat cells, a CD4 positive lymphoma cell line (53). After treatment with 1 μM of VT107, Jurkat cells showed enhanced cytotoxicity towards co-cultured 4T1 cells (FigS4C). Jurkat cells’ cytotoxicity was also significantly increased in shTEAD-transduced lines (targeted at TEAD 1,3 & 4 Fig S4D) compared to control (Fig 4G). RNA-seq analysis of these TEAD knockdown Jurkat lines revealed a significant upregulation of T cell survival and activation markers such as CD28 and CD86 (54,55) and cytotoxicity granzymes K and M as well as Th1 cell differentiation markers CCR5 and EOMES (56,57) (Fig 4H). Signaling pathway analysis further confirmed Th1 pathway enrichment and immune activation and downregulation of T cell exhaustion signaling pathway and IL10 signaling (58) (Fig 4I). Taken together this data supports a direct effect of TEAD-controlled pathways on T cell functions, activation of Th1 cells and CD8+ T cell cytotoxicity.

### TEAD inhibition affects T cell-macrophage crosstalk and enhances cancer cell elimination

To investigate whether the dual effect of iTEAD treatment on myeloid and T cells can enhance their crosstalk in the lung microenvironment, we first isolated and characterized peritoneal macrophages +/− iTEAD (Fig. 5A). We observed that treatment with VT107 decreased gene expression of M2-like and fibrosis regulators, Aldh1a1 and Ccl17 (59), as well as TEAD4 and its downstream target gene Ankrd1. Pvalb, an inhibitor of M2-like phenotype, was the top upregulated gene (60) (Fig. S5A). Overall pathway analysis showed downregulation of disease-related chronic inflammation while cell cycle related pathways were upregulated (Fig. S5B). The iTEAD pretreated macrophages and T cells isolated from the spleen of tumor naïve mice (Fig 5A) were then mixed for 5 hours before collecting the conditioned media for analysis. IL12 and IL6 levels were significantly increased in the iTEAD-treated cell mix (Fig. 5B). No significant changes were detected in the other factors on the array. Next, 4T1 cells were plated and allowed to attach on electrode-coated arrays to monitor cancer cell viability after the addition of macrophages +/− iTEAD and T cells (Fig 5A). VT107-treated macrophages significantly increased untreated T cells’ cytotoxicity compared to T cells alone and this crosstalk was diminished by neutralizing antibodies for IL12 (Fig 5C). This reversal of T cell cytotoxicity by IL12 antibodies was not observed in VT107-treated T cells in the absence of macrophages (Fig S5C) indicating a critical role for IL12 in the crosstalk between macrophages and T cells. In 3D, 4T1 spheres that were embedded into a collagen I and matrigel mix, showed elevated signal of cleaved caspase-3, a marker of cellular apoptosis, in the presence iTEAD-treated macrophages and T cells (Fig. 5D and 5E) This effect was again reversed by an IL12 neutralizing antibody. The increase in cleaved caspase-3 in 4T1 cells in the presence of VT107-treated T cells, was not detected at 24 hours in the absence of macrophages (Fig S5E). However, it can be seen at later time points (e.g in Fig. 4D after 72 hours iTEAD treatment). Overall, the data supports a model where iTEAD treatment increases the macrophage-T cell crosstalk through IL12. The iTEAD impact is further enhanced by increasing direct T cell cytotoxicity mediated through IL-2 (Fig 4).

**Figure 5:**
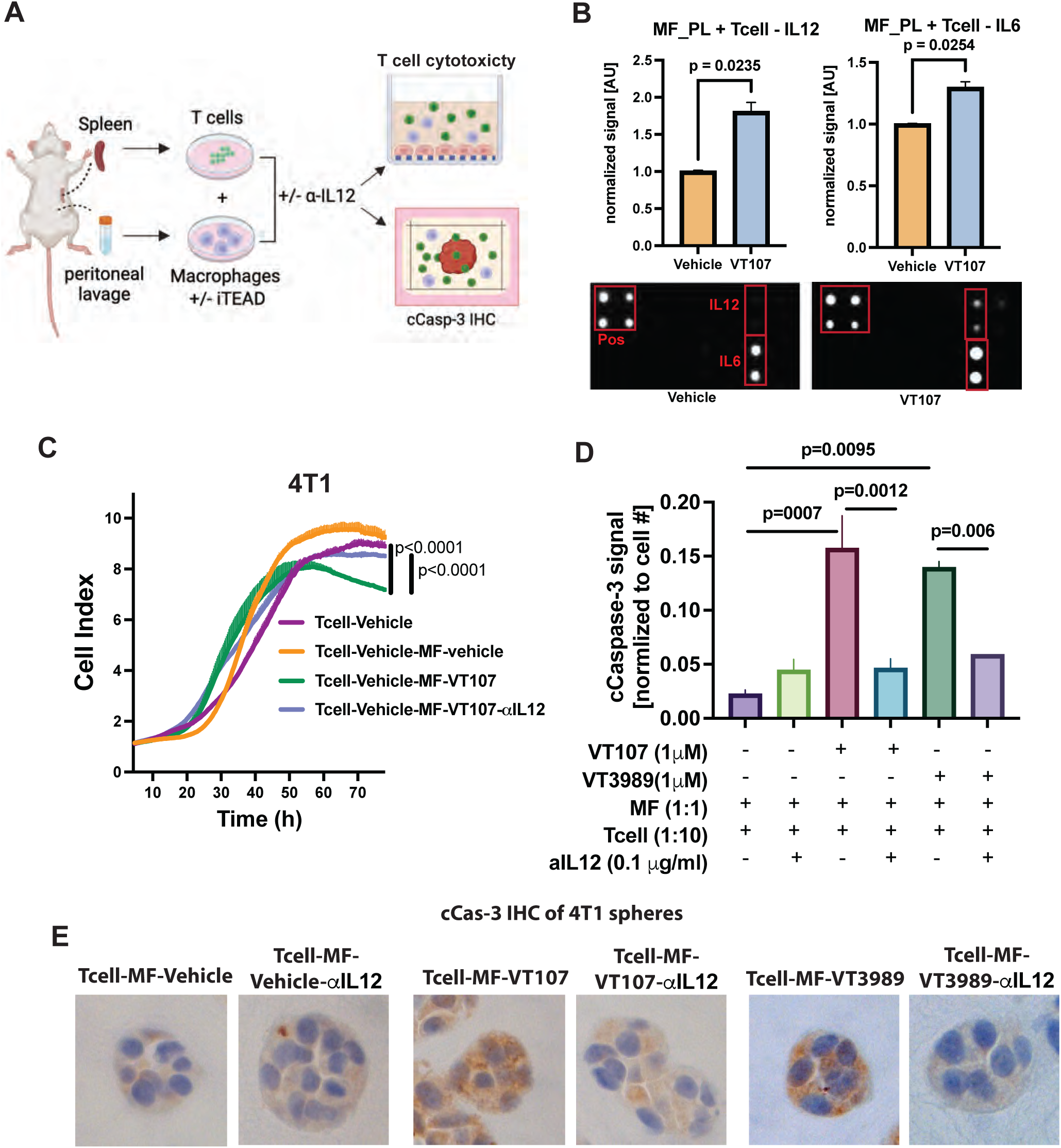
TEAD inhibition affects T cell-macrophage crosstalk and enhances cancer cell apoptosis. A) Schematic of experiments macrophage and T cell isolation and downstream crosstalk analysis. T cells were isolated from naïve mice spleens. Macrophages were isolated from the peritoneal cavity by lavage (MF-PL) and pretreated for 24hrs with 1μM VT107. B) Blot of cytokine array and quantified data showing changes in secreted factors in the conditioned media of T cells and macrophages mixed for 5 hours. (Mean ± SEM, student t-test). C) RTCA of 4T1 cancer cells measured by electric impedance sensing of attached cells (cell index) over time. IL12 neutralizing antibody (0.1 mg/ml) was added to the macrophages as indicated. (Mean±SD, student t-test). D) Quantification of the cCas-3 IHC signal 24 hours after adding the immune cells to the 4T1 spheres. n=3-10 spheres per condition. (Mean±SD, student t-test). E) Representative images of immunohistochemistry of cCas-3 on 4T1 spheres embedded in Matrigel/ collagen mix. T cell and peritoneal macrophages were added to the spheres with the indicated treatments (vehicle, 1μM VT107,VT3989, or 0.1 mg/ml IL12 antibody).

## Discussion

In this study, using TEAD inhibitors we have revealed a new aspect of TEAD signaling in the lung immune stroma that enhances the anti-metastatic effects of these agents. We found that TEAD-regulated transcription alters the immune microenvironment during metastatic progression in the lung. The tumor promoting macrophages were decreased along with pro-tumor inflammation pathways. TEAD inhibitor treatment increased the number of cytotoxic CD8 T-cells and Th1 CD4+ T-cells present within the lungs and enhanced the crosstalk between macrophages and T cells via IL12 (Fig 6). We also showed that these parameters were not affected in the primary tumor suggesting a tissue specific effect on resident immune cells in the lungs. In fact, the lung resident alveolar macrophage population increased and shifted away from an M2-like phenotype in the iTEAD-treated lungs. In a recent study, an increase in alveolar macrophages was observed in the lungs of short form Ron knockout mice, a receptor expressed by tissue resident macrophages that exhibited a significant reduction in lung metastasis (61). Another study showed that alveolar macrophages act as an innate immune barrier to metastatic outgrowth of dormant cancer cells that have seeded in the lungs of a breast cancer mouse model (62). These data support an important role for resident alveolar macrophages in halting the progression of metastatic lesions in the lung.

**Figure 6:**
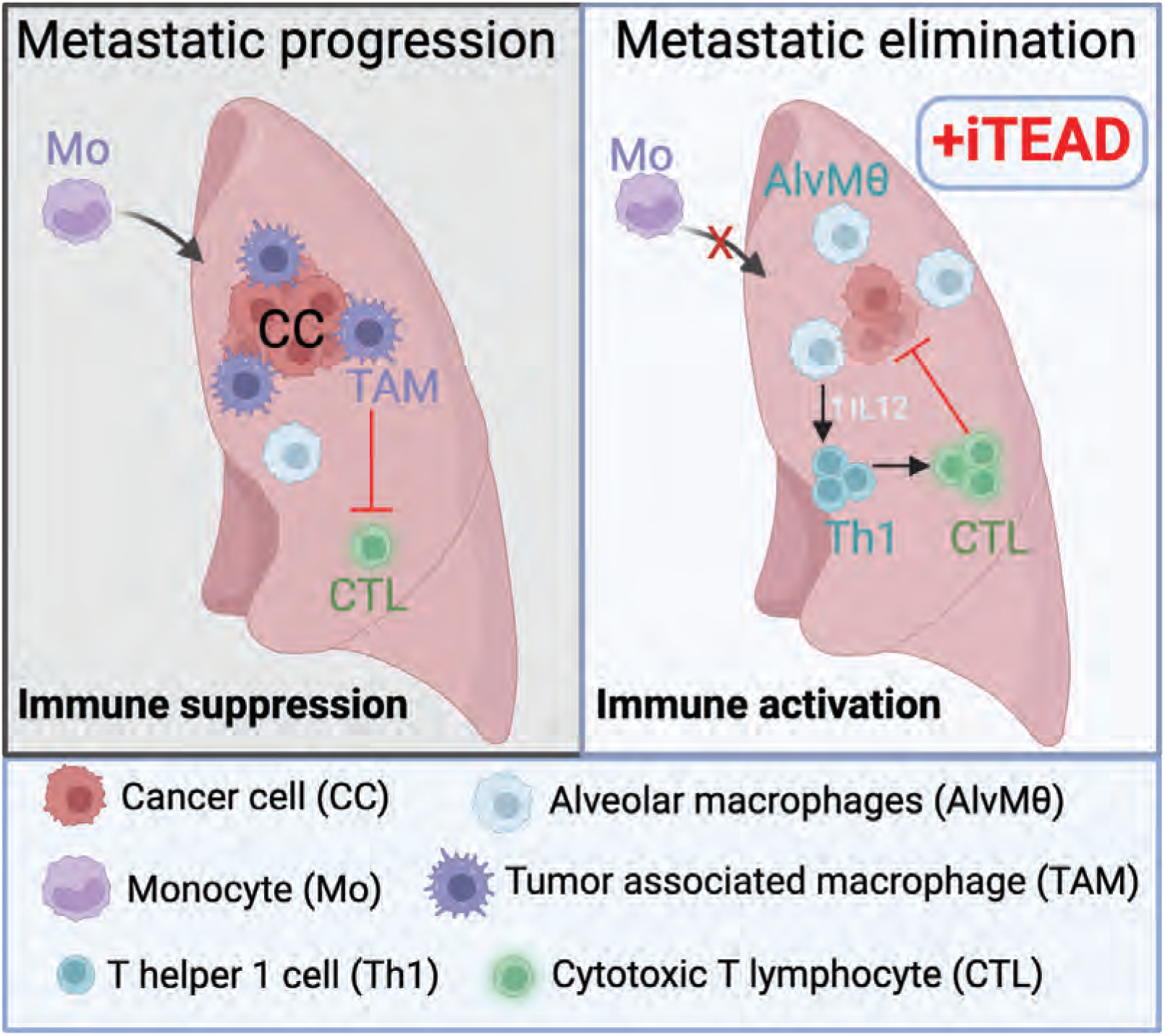
Left panel: During metastatic progression, bone marrow-derived monocytes (Mo) are recruited to the lung and tumor associated macrophages (TAM) are reprogramed to support tumor growth and immune suppression by inhibiting cytotoxic T lymphocytes (CTL). Right panel: In iTEAD treated lungs, Mo and TAMs are reduced while anti-tumor alveolar macrophages (AlvMø) increase along with IL12 signaling. This is accompanied with an increase in T helper 1 cells (Th1) and cytotoxic T lymphocytes (CTL) to attain immune activation necessary for elimination of poised metastases.

Primary tumors in breast cancer preclinical models showed no response to TEAD inhibitor treatments as reported here in this study (Fig.1B and Fig 3B) and by communication with scientists from Vivace therapeutics (19). However, our data reveals a distinctive mechanism aimed at reprogramming the immune microenvironment in the metastatic niche of the lungs. Here we showed that TEAD inhibitor treatment has changed the phenotype of multiple immune cells and their interactions. TEAD palmitoylation inhibitors are now in multiple clinical trials due to the promising results in preclinical models and low toxicity (63). They were originally tested on neurofibromatosis type 2 (NF2) negative or mutant cell lines, a tumor suppressor and upstream regulator of the Hippo pathway encoding the Merlin protein and showed anti-proliferative effects. There was no impact in NF2 wildtype, Merlin-positive mesothelioma cell lines Furthermore, TEAD inhibitors, including VT107, showed anti-proliferation in mesothelioma cell lines that had no detected NF2 mutations but are Merlin-negative by Western Blotting analysis (32) Initial results from VT3989 clinical trial showed a benefit in patients with metastatic tumors regardless of their NF2 status (17) suggesting that NF2 mutation status of the primary tumor is not indicative of the anti-metastatic effect of TEAD inhibitors. Merlin protein analysis by IHC is needed to determine the Merlin expression of these tumors. Here we showed that TEAD inhibition had no effect on the primary tumor in two different mouse models that are NF2 wildtype, yet there was a significant reduction in lung metastasis. Pathway analysis in different immune cell lines showed the various roles of TEAD transcription in a cell type dependent manner. While TEAD inhibition reduced proliferation of cancer cells that are NF2-mutant, it had an activating effect on T cells, upregulating survival and activation markers such as CD28 and CD86, as well as cytotoxic granzyme K and M (Fig. 4). TEAD inhibition had a differential effect on secreted IL12 levels in lung versus bone-marrow derived macrophages discerning a cell type specific effect (Fig.2 E and F). The TEAD family of transcription factors has four members that are differentially expressed in tissue types (64,65), and this may also contribute to the diverse effects of TEAD inhibitors. Further studies delineating the role of specific TEADs in the different tissue types is still needed.

The YAP/TEAD complex has been reported to induce inflammation that promotes cancer growth, invasion, and survival (26,66,67). We showed that TEAD inhibition repressed inflammatory pathways associated with tumor progression in the lung tissue and reversed the tumor associated macrophages’ phenotype that supports immune suppression and evasion (Fig. 3) (68). T cell exhaustion and exclusion is largely due to the suppressive signals of chronic inflammation in the tumor microenvironment (69). However, TEAD inhibition fostered an acute phase inflammation profile and enhanced macrophage-T cell interaction via IL12 signaling (Fig 2 and Fig 5). Due to the large evidence highlighting the role of macrophages in reshaping the tumor microenvironment (44), novel therapeutics are being aimed at reprogramming TAMs or inhibiting TAMs associated secretome, particularly by IL12 (70). Systemic administration of IL12 showed tumor response and immune re-activation (71,72), but did not perform well in clinical trials due to the toxicity associated with the high levels of systemic IL12 (73,74). Thus, other approaches that can induce physiologic levels of IL12 may have a better success rate in the clinic. For instance, a study in ovarian cancer showed a reduction in tumor size and metastasis upon treatment with monophosphoryl lipid A (MPLA) and interferon γ (IFNγ) converting protumor CD206+ TAMs to inducible NO synthase (iNOS)+ macrophages that activated cytotoxic T-cells through macrophage-secreted IL12 (75). Genetically engineered myeloid cells that delivered IL12 to metastatic sites, but were unable to expand *in vivo,* reversed immune suppression and activated anti-tumor immunity (34). Our data established a novel role of TEAD inhibitors as modulators of the inflammatory profile in the lungs, by reprogramming the macrophage phenotype and enhancing macrophage and T cell crosstalk via IL12 signaling (Fig 6). Consistent with published reports that emphasize IL12 signaling in anti-tumor immune response, the upregulation of IL12 production and signaling pathway upon iTEAD treatment have led to the reversal of the immunosuppressive program that is initiated by metastatic cells at the new microenvironment.

The fact that TEAD inhibitors alter the immune microenvironment that impacts the outgrowth of poised metastasis, supports the broad use of TEAD inhibitors on other metastatic cancers. Further studies to determine the role of TEAD in the immune milieu of different tissues and other metastatic niches such as the liver, brain and bone are warranted. Furthermore, combination of TEAD inhibitors with immunotherapy for metastatic disease may be a promising therapy regimen. Our results shed light on the direct effect of iTEAD in T cell activation and survival and importantly the sequential events related to the interaction between innate and adaptive immune cells. iTEAD-treated macrophages were able to activate T cell cytotoxicity within 24 hours on tumorspheres (Fig.5D and 5E). However, T cell cytotoxicity directly activated by iTEAD treatment was only observed after 72 hours (Fig. 4D). This data could inform our decision on which immunotherapy would best synergize with TEAD inhibitors. The mechanism of action for CTLA-4 and PD-1 is distinct; CTLA-4 blockade is important in the early priming of T cells and it’s been implicated in the differentiation of CD4+ T cells particularly Th1 cells. PD-1 antibodies impact late stages of T cell activation that are PD-1+ and does not influence T cell differentiation (76–78). Thus, TEAD inhibitors and CTLA-4 antibodies may better synergize in their impact on Th1 CD4+ T cells and the initial stages of T cell activation.

Using TEAD inhibitors, we delineated a new role for TEAD mediated pathways that influence resident macrophage function, T cell activation, and their crosstalk to create an immune activated microenvironment that suppresses metastatic outgrowth in the lungs. We have provided valuable insights into the changes in the tumor immune cell populations impacted by TEAD-regulated transcription and how its inhibition can be harnessed to activate an antitumor immune microenvironment. Our results provide a rationale for using TEAD inhibitors to enhance immunotherapy in treating metastatic breast cancer which is limited by protumor inflammation and immune suppression (79).

## Methods

### Real-Time Cell Analysis

Cell analysis was done using E-plates from xCELLigence, (Agilent, # 5469830001) according to the manufacturer’s protocol. For cell cytotoxicity, cancer cells (10,000 – 25,000 cells per well) were seeded and allowed to attach for 1-2 hours prior to the addition of immune cells and/or drug treatments. For cell proliferation, cancer cells were seeded at 5,000-10,000 cells per well with incremental doses of drug treatments or DMSO as a vehicle control. Cell index was measured by changes in electric impedance at 15min intervals. TEAD inhibitors were obtained from Vivace therapeutics.

### Histology

Formalin-fixed, paraffin embedded (FFPE) lung tissue sections were stained with hematoxylin and eosin. Immunohistochemistry staining was performed on FFPE tumor and lung tissue sections. Slides were deparaffinized in xylene and rehydrated following a series of ethanol washes. Heat-induced antigen retrieval was then performed by placing slides in warmed sodium citrate buffer (pH 6.0) (Invitrogen, 005000) in a pre-heated steamer for 20 minutes. Slides were cooled at room temperature for an additional 20 minutes, prior to quenching endogenous peroxidase activity using 3% hydrogen peroxide buffer (Fisher, BP2633500) for 10 minutes. Next, slides were incubated overnight at 4°C with either anti-CD8 (Cell Signaling Technology, #98941), anti-CD68 (Abcam, ab303565), anti-CD206 (Cell Signaling Technology, #24595), or anti-F480 (Cell Signaling Technology, #70076). Following PBS washes, the slides are then stained with a biotinylated secondary antibody using the VECTASTAIN Elite ABC-HRP Kit according to the user’s manual (Vector Laboratories, PK-6101). Positive brown staining was produced by applying ImmPACT DAB EqV Substrate Kit (Vector Laboratories, SK-4103) to the slides, which were then counterstained with hematoxylin (Sigma, MHS16). Images were captured using the Olympus BX40 microscope, and positive staining was quantified manually. Immunofluorescent staining was performed on lung tissue sections. Slides were baked at 60°, deparaffinized in xylene, rehydrated, washed in DI water and incubated with 10% neutral buffered formalin (NBF) for an additional 20 minutes to increase tissue-slide retention. Epitope retrieval/microwave treatment (MWT) for all antibodies was performed by boiling slides in Antigen Retrieval buffer 6 (AR6 pH6, Akoya AR6001KT). Protein blocking is performed using antibody diluent/blocking buffer (Akoya, ARD1001EA) for 10 minutes at room temperature. Slides were incubated with Luciferase antibody (Sigma, #L0159) for 1 hour and OPAL Fluoresce 620 then counterstained with spectral DAPI (Akoya FP1490) for 5 min and mounted with ProLong Diamond Antifade (ThermoFisher, P36961). Slides were scanned at 10X magnification using the Vectra 3.0 Automated Quantitative Pathology Imaging System (PerkinElmer/Akoya,).

### Western blot

Cells were lysed in NP40 lysis buffer, in the presence of cOmplete Protease Inhibitor Cocktail (Roche, #12352204), and a phosphatase inhibitor Na_3_VO_4_. Lysates were separated on an SDS-PAGE gel and immunoblotted with antibodies against pan-TEAD (Cell Signaling Technology, #13295), and GAPDH (Cell Signaling Technology, #2118).

### T-cell Isolation from tissues

Mouse lungs and tumors were digested using the Tumor Dissociation Kit (Miltenyi Biotec, #130-095-929) as described in the manufacturer’s protocol. Tumors and lungs were subdued to mechanical dissociation for 60 and 30 minutes respectively. Spleens were passed through a 100um cell strainer (Fisherbrand, #22-363-549) using a syringe plunger. T-cells were isolated from all dissociated tissues using the EasySep Mouse T-Cell Isolation Kit (StemCell Technologies, #19851).

### Bone marrow isolation and macrophage differentiation

The bone marrow was flushed with 100ul phosphate buffered saline (PBS) from the femur and tibia of a BALB/c mouse, treated with ACK lysis buffer (Thermofisher, A1049201) and cultured in RPMI medium with 10% FBS. Cells were cultured with 10ng/ml mouse M-CSF (StemCell, 78059.1) for 3 days then washed with PBS to get rid of floating cells. Adherent cells (macrophages) were treated with vehicle or 1 μM VT107 for 24 hours.

### Flow cytometry

Single cell suspensions were washed 2x with PBS. Cells were then stained with Zombie NIR Fixable Viability Kit (Biolegend, 423106) or Zombie Aqua Fixable Viability Kit (Biolegend, 423102) for 20 minutes at room temperature, to identify dead cells. Next, cells were washed with FACS buffer (1X PBS + 2% FBS) and incubated with conjugated antibodies binding to cell surface proteins, at 4°C for 30 minutes. A combination of the following antibodies were used: anti-CD45-Spark Blue 550 (Biolegend, 103166), anti-CD3-PercP/Cy5.5 (Biolegend, 100217), anti-CD4-Pacific Blue (Biolegend, 116007), anti-CD8-BV785 (Biolegend, 100750), anti-CD25-BV650 (Biolegend, 102038), anti-CD45-AF700 (Biolegend, 103128), anti-CD45R (B220) (Biolegend, 563103), anti-SiglecF-APC (Miltenyi Biotec, 130-102-241), anti-CD11b-PE/Dazzle (Biolegend, 108745), anti-CD11c-FITC (Biolegend, 117306), anti-F480-PE/Cy7 (Biolegend, 123114), anti-Ly6G-BV785 (Biolegend, 127645), anti-Ly6C-BV570 (Biolegend, 128030), anti-CD206-AF700 (Biolegend, 141734), and anti-CD86-AF488 (Biolegend, 105018). Cells were fixed and permeabilized using the BD Cytofix/Cytoperm Fixation/Permeabilization Kit (BD Biosciences, #554714) prior to incubation with anti-Foxp3-AF647 (Biolegend, 126407), anti-Gata3-PE (Biolegend, 653803), anti-RORγ t-BV421 (Biolegend, 562894), and anti-T-bet-BV711 (BD Biosceinces, 4B10). After staining, cells were washed 2x and subjected to flow cytometry analysis using the BD FACSymphony A3 flow cytometer analyzer. Results were analyzed using FCS Express 7 Research (De Novo Software).

### Degranulation assay of T-cells

T-cells were isolated from the spleens of naïve BALB/c as described above and treated *in vitro* with DMSO control or VT107 for 24 hours. T-cells were then co-cultured with 4T1 cells with the same treatment conditions. After 72 hours, all cells were collected and subdued to flow cytometry using anti-CD45-AF700 (Biolegend, 103128), anti-CD8-BV785 (Biolegend, 100750), anti-CD4-Pacific Blue (Biolegend, 116007), anti-CD107a-PE/Cy7 (Biolegend, 121620), and Helix NP Blue (Biolegend, 425305). Analysis was performed on the BD FACSymphony A3 flow cytometer analyzer, and results were analyzed using FCS Express 7 Research (De Novo Software).

### Cytokine Array

Cells were grown in their recommended cell culture media and treated with 1μM VT107 for 24 or 72 hours. Conditioned media was collected and passed through a 0.2 micron PES filter then analyzed on a RayBio® Mouse Cytokine Array (#AAM-TCR-1) according to the manufacture’s protocol. The array includes two spots for each cytokine. The luminescence signal was normalized to the positive control on each membrane, then to the vehicle treatment signal of each cytokine.

### Animal experiments

Studies in mice were reviewed and approved by the Georgetown University Animal Care and Use Committee (GUACUC). Animals were randomized to receive vehicle control (DMSO) or VT107. Treatment was formulated in 5% DMSO, 10% solutol and 85% D5W; D5W. 5% glucose and administered daily by oral gavage. Five hundred thousand cells were injected into the mammary fat pad of age-matched 8 weeks old female NOD/SCID or C57/Bl6 mice purchased from Charles River or Jackson Laboratory respectively. A hundred thousand 4T1 cells were injected into the tail veins of BALB/c mice purchased from Jackson Laboratory. Experimental timelines, treatment start times and durations are indicated in the figure diagrams and figure legends. Tissues collected were preserved in RNAlater (Thermo fisher, # AM7020) or fixed in 10% formalin for histology analysis.

### RNA sequencing

The total RNA from tissue or cell lines was extracted using RNeasy Mini Kit (Qiagen, # 74104) according to the manufacturer’s instructions. Total RNA integrity was assessed with the Agilent 2100 Bioanalyzer. The library preparation and next-generation sequencing (NGS) were performed at Novogene Corporation Inc. (Sacramento, CA, USA). At least triplicate samples per experimental condition were analyzed, raw sequence reads (150bp paired end, >45×106 average reads/sample) were aligned to the human or mouse transcriptome using STAR, and aligned reads translated to expression counts via featurecounts, followed by a standard edgeR (RRID:SCR_012802) pipeline to identify DEGs under specific conditions. Ingenuity pathway analysis (IPA, RRID:SCR_008653) was undertaken using a list of all differentially expressed genes with a cutoff of p-value less than 0.05 to identify signaling pathways and upstream regulators.

### Cell lines and shRNA

MCFDCIS (RRID:CVCL_5552) cell line was maintained in DMEM/F12 (1:1) medium (Gibco, Waltham, MA, USA, 11039-021) with 5% horse serum, 20 μg/mL epidermal growth factor (EGF), 100 μg/mL hydrocortisone, 10 μg/mL insulin, and 100 ng/mL cholera toxin. E0771 (RRID: CVCL_GR23) cell line was maintained in DMEM medium (Gibco, 11995-065) with 10% fetal bovine serum (FBS). 4T1 (RRID: CVCL_0125), THP1 (RRID: CVCL_0006) and Jurkat (RRID: CVCL_0065) cell lines were maintained in RPMI medium with 10% FBS. All cells were cultured at 37 ◦C with 5% CO2. Lentivirus was made by transfecting HEK293T cells with packaging and envelope plasmids and shRNA plasmids. Media containing virus was collected 48, filtered and virus particles pelleted with PEG-it (System Biosciences, #LV810A-1) according to manufacturer’s instructions. Cells were infected with lentivirus then selected with 5mg/ml puromycin (Thermofisher, #A11138-03).

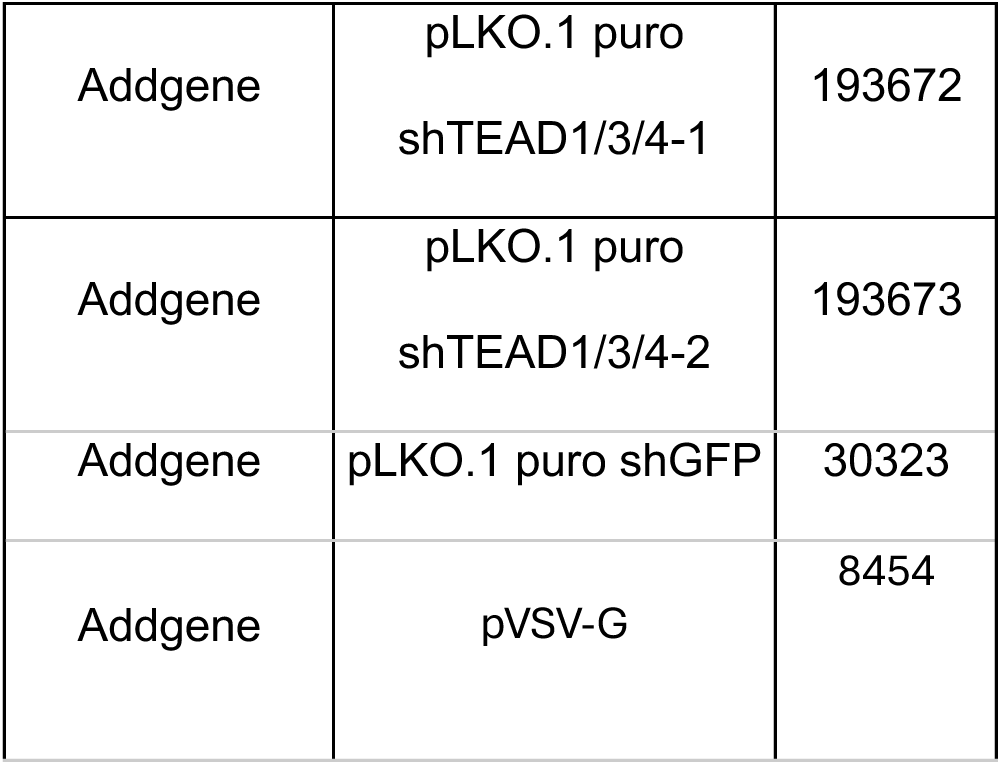
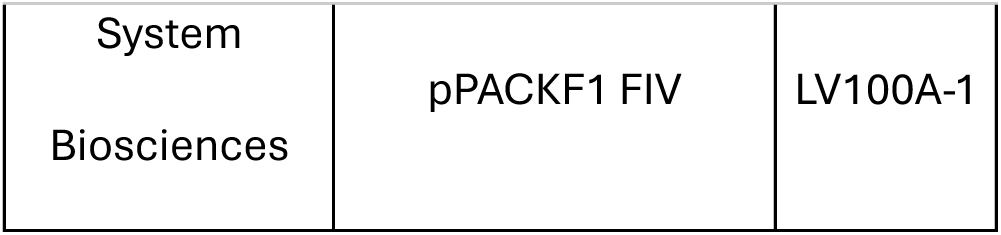

### RNA extraction and qPCR

Total RNA was extracted from cell lines using a RNeasy Mini Kit (Qiagen, # 74104) according to the manufacturer’s instructions. cDNA from total RNA was made with the iScript cDNA synthesis kit according to the manufacturer’s protocol (Biorad, # 170-8891) and qPCR was performed in an iCycler iQ (BioRad) using the iQ SYBR Green Supermix (BioRad, # 170-8882). Primers used:

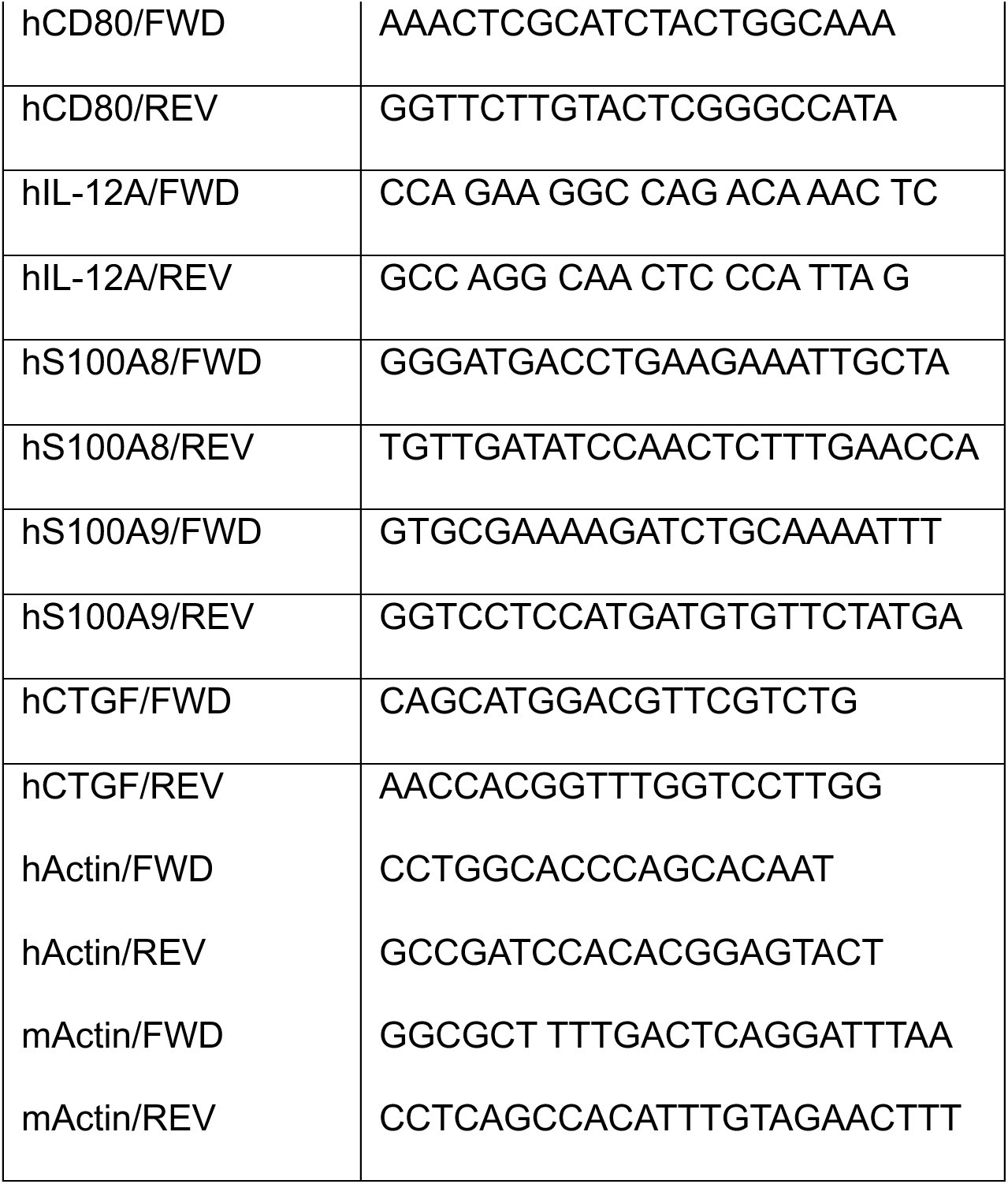

### 3D sphere co-culture assay

One thousand 4T1 cells were aggregated in 81-well agarose molds (Microtissues, #Z764019) Immune cells and treatments were added to the media according to the experiment’s description. At the endpoint, spheres were fixed in 5% formalin and embedded in histogel (Fisher Scientific, #22-110-678) for sectioning. FFPE immunohistochemistry was done as previously described in this methods section with primary antibodies for Caspase 3(1/80, BIOCARE, CP229A). Images were captured using the Olympus BX40 microscope, and DAB signal per sphere was quantified in imageJ and normalized to the number of nuclei.

In the experiments where spheres were embedded in 20% Matrigel (Corning, #354230) and 80% collagen I mix (Thermofisher, #154453), agarose molds were removed before immune cells and treatments were added in the media and onto the matrix embedded spheres according to the experiment’s description. THP1 cells were labeled with 1ul of the CMFDA green tracer (Thermofisher, C7025) for 30 min, then washed with RPMI media.

### Statistics

Analyses were performed either using the R platform for statistical computing (version 3.6.1) and the indicated library packages implemented in Bioconductor (RRID:SCR_006442) or Prism 7 (Graphpad Inc, RRID:SCR_006442). Student t-tests and Fisher exact tests were used for comparisons as indicated in figure legends, with p<0.05 as the threshold for statistical significance in all tests.

## Data Availability

The RNA-sequencing data have been deposited to the Gene Expression Omnibus under record GSE290381

## Acknowledgments

We thank Dr. Allejandro Villagra and Dr. Tracy Tang for reviewing the manuscript. We thank Vivace Therapeutics for providing the TEAD inhibitors (VT107 and VT3989). We thank Lombardi shared resource facilities; the Tissue Culture and Biobanking, Flow Cytometry and Cell Sorting, Microscopy and Imaging, Histopathology and Tissue, and Animal Models.

## Authors contributions

GMS conceived and directed the study. RR, MOS, MM, TA, RB, AK and GMS designed and performed experiments and analyzed the data. GS and RR wrote the manuscript. MOS, MM, TA, RB, AK, AW and ATR provided input and reviewed the manuscript.

---

**Figure S1.**
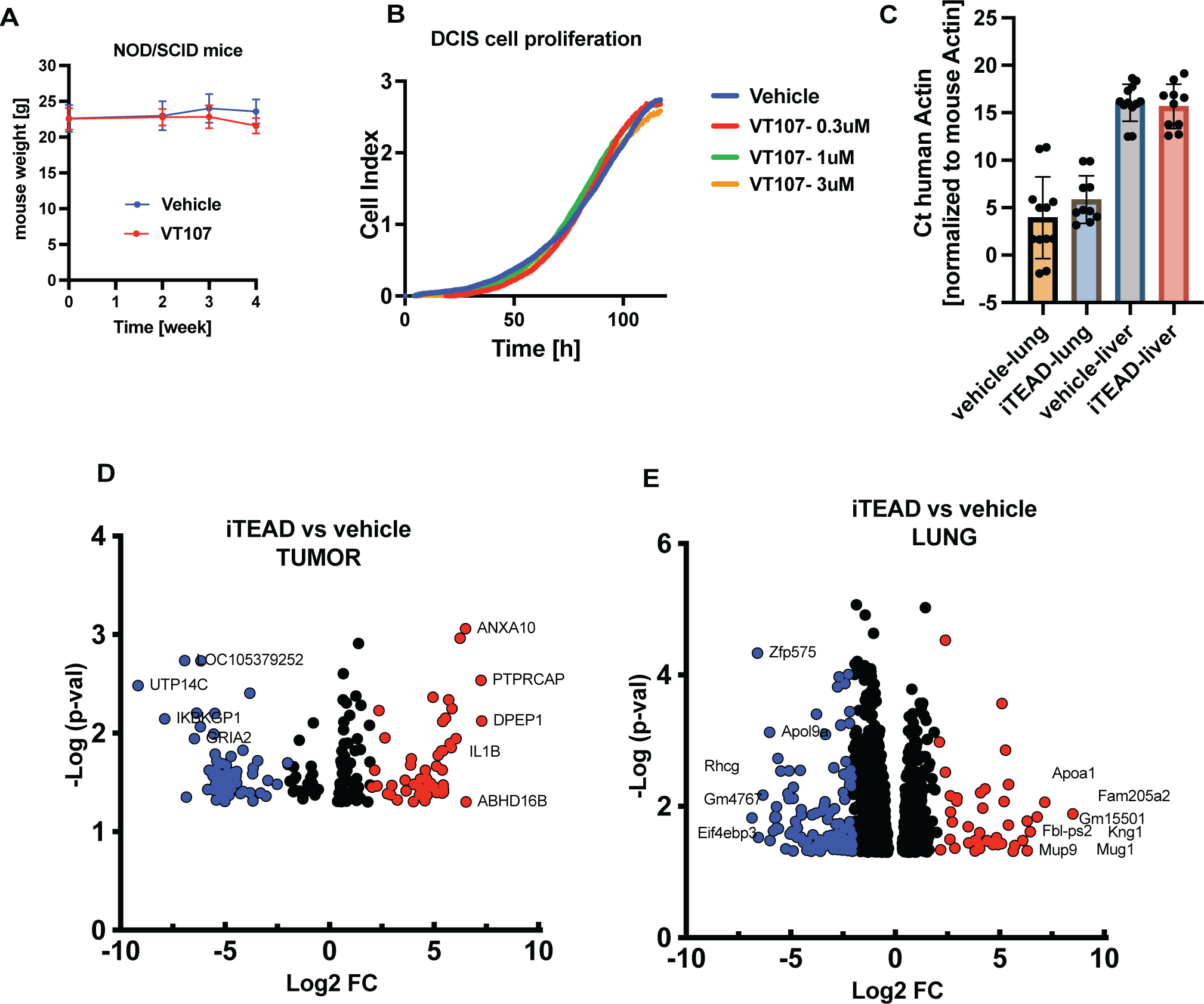
A) The body weight of NOD/SCID mice bearing DCIS:DCISΔ4 tumors in the different treatment groups. B) Real-Time Cellular Analysis (RTCA) of proliferation of DCIS cells measured by electric impedance sensing of attached cells (cell index) over time with increasing doses of VT107. (Mean±SD, student t-test). C) qPCR of human actin (cancer cells) normalized to mouse actin (stroma) in the lung (metastasis site) and liver (control organ) tissue of mice that were treated with VT107 or vehicle. D) Volcano plot depicting change in gene expression in tumor tissues in response to VT107 treatment of NOD/SCID mice. E) Volcano plot depicting change in gene expression in lung tissues in response to VT107 treatment of NOD/SCID mice. RNA-sequencing reads from tumors were aligned to the human transcriptome while lung tissue reads were aligned to the mouse transcriptome. Red and blue circles indicate regulated genes with −Log p-value >1.3 and |log_2_FC|=2. A full list of changed genes is in supplementary table 1.

**Figure S2:**
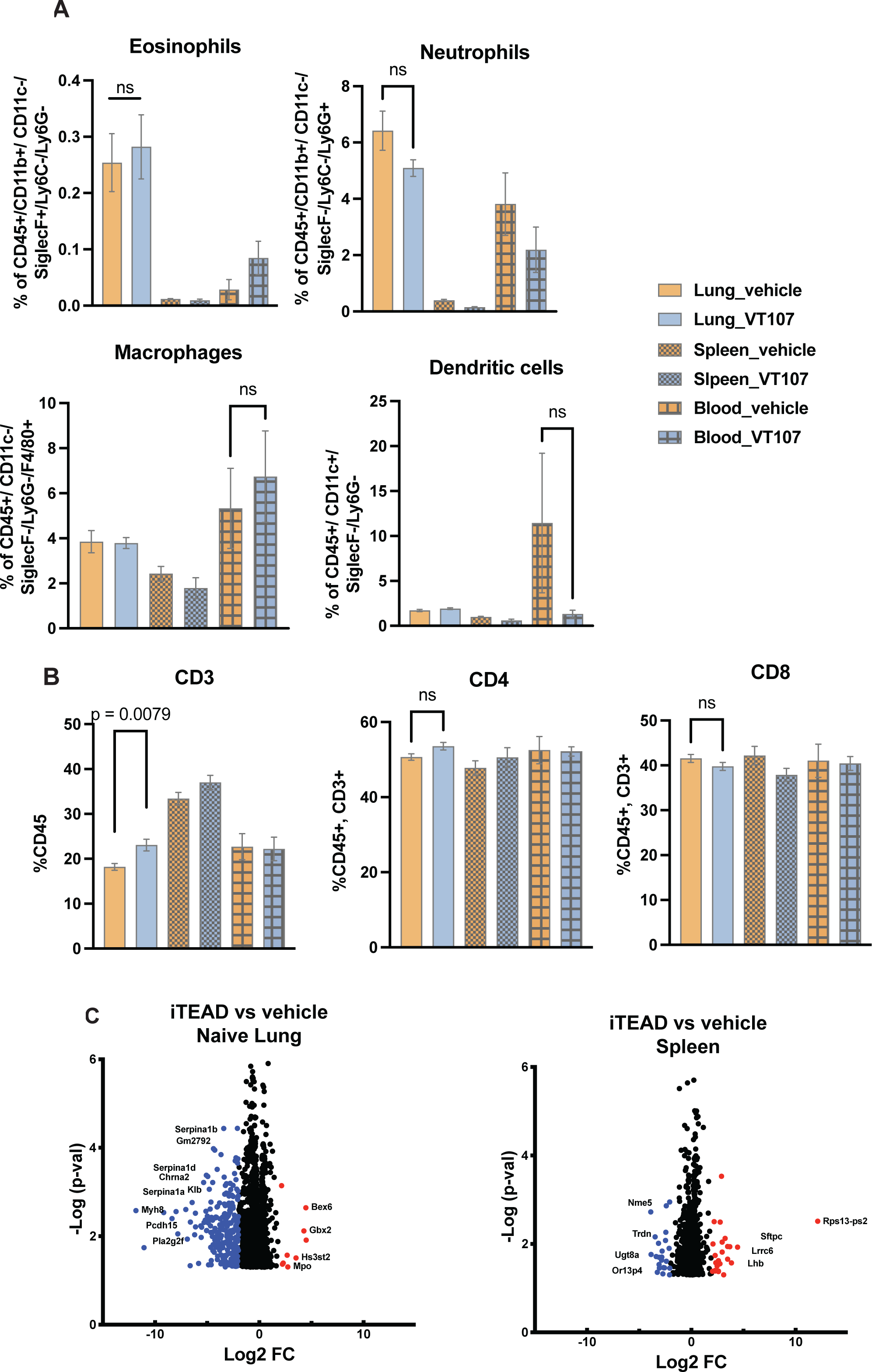
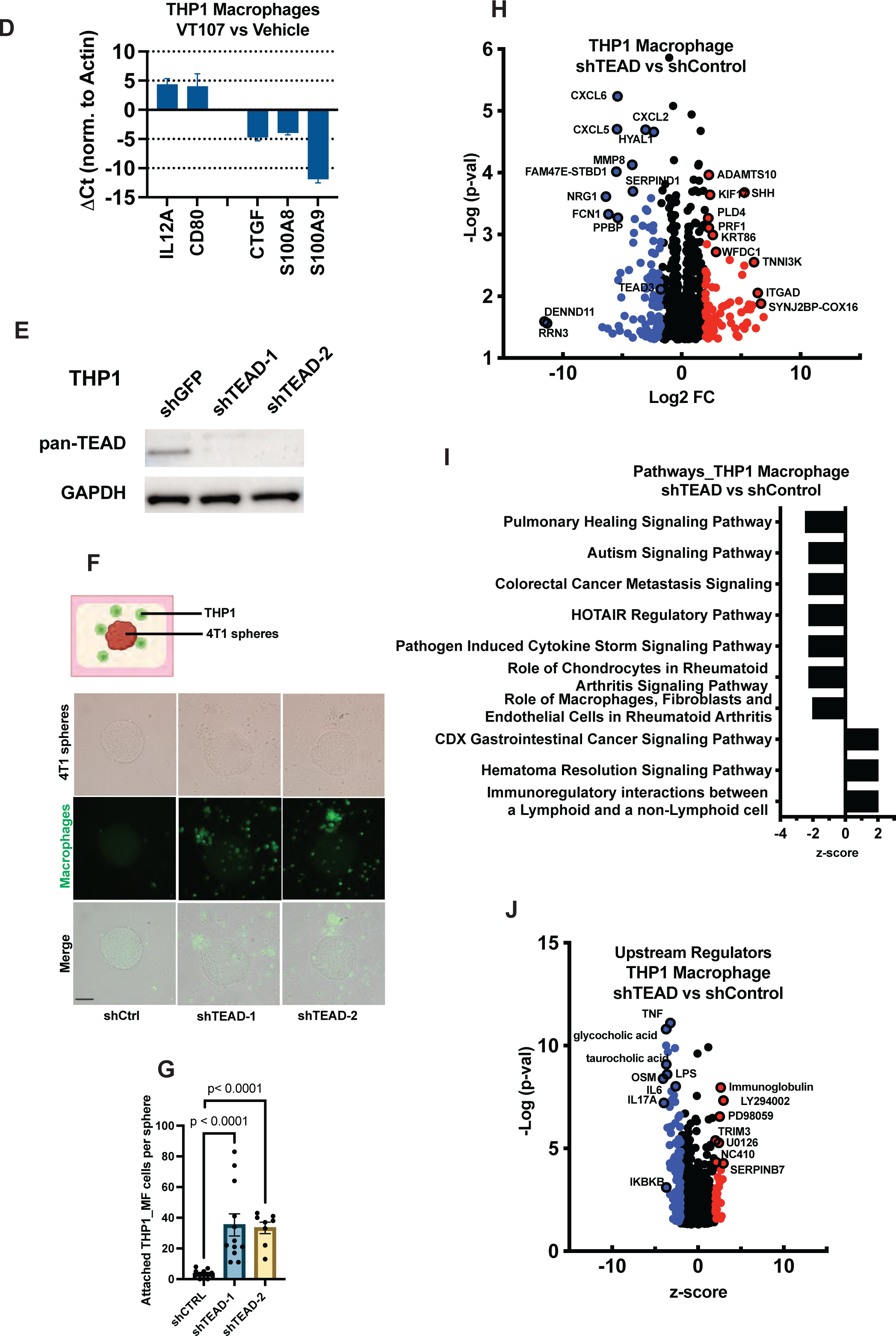
A) Flow cytometry analysis showing change in the eosinophils, neutrophil, macrophages or dendritic cell populations in naïve mice treated with VT107 or vehicle from Fig 2A. B) Flow cytometry analysis showing change in lymphoid cells of naïve mice treated with VT107 or vehicle from Fig 2A. C) Volcano plots depicting changes in lung and spleen gene expression between VT107 or vehicle treatment from Fig 2A. Red and blue circles indicate regulated genes with −Log p-value >1.3 and |log_2_FC|=2. A full list of changed genes is in supplementary table 1. D) Changes in gene expression were measured by qPCR in THP1 macrophages that were treated with 1μM VT107 or vehicle. The gene expression of three TEAD target genes is downregulated confirming TEAD-related transcription inhibition by VT107. E) Western blot of pan-TEAD protein levels in THP1 monocytes infected with either control shRNA or two distinct shTEAD lentiviruses. GAPDH protein levels were used as a loading control. F) Schematic of THP1 macrophages co-cultured with 4T1 spheres embedded into a collagen I: Matrigel mix.THP1 macrophages harboring two different shRNAs for TEAD or control were stained with a green cell tracer. G) Quantification of THP1 macrophages that invaded into the matrix and attached to the 4T1 spheres. n=8-12 spheres per condition. (Mean±SD, student t-test). H) Volcano plots depicting changes in gene expression in THP1 macrophages between shTEAD and shControl. Red and blue circles indicate regulated genes with −Log p-value >1.3 and |log_2_FC|=2. A full list of changed genes is in supplementary table 2. I) IPA showing altered signaling pathways in shTEAD versus shControl THP1 macrophages with p-val<0.05 and a |z-score| > 2. J) IPA showing upstream regulators altered in shTEAD versus shControl THP1 macrophages with a −Log p-value >1.3 and a |z-score| > 2.

**Figure S3.**
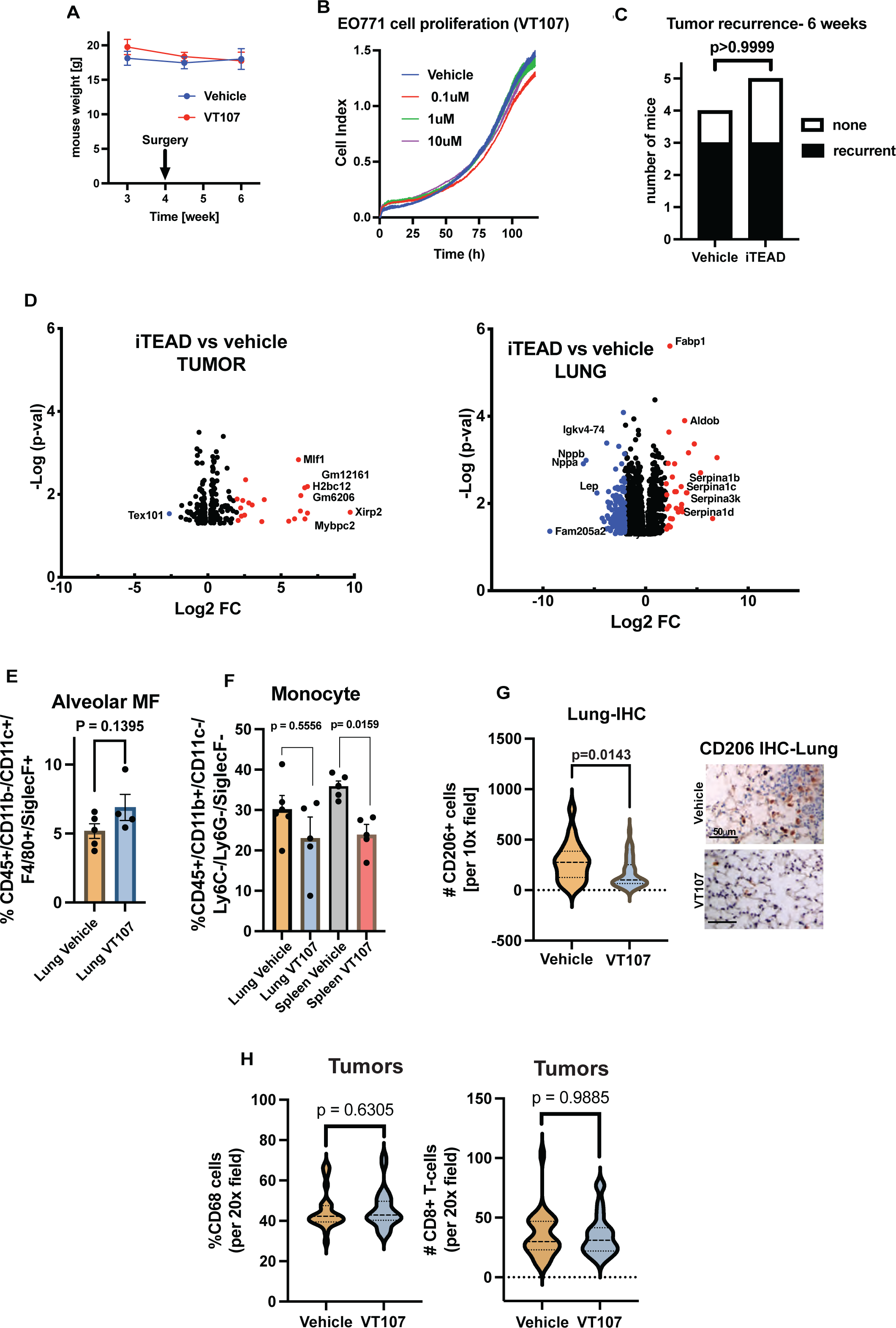
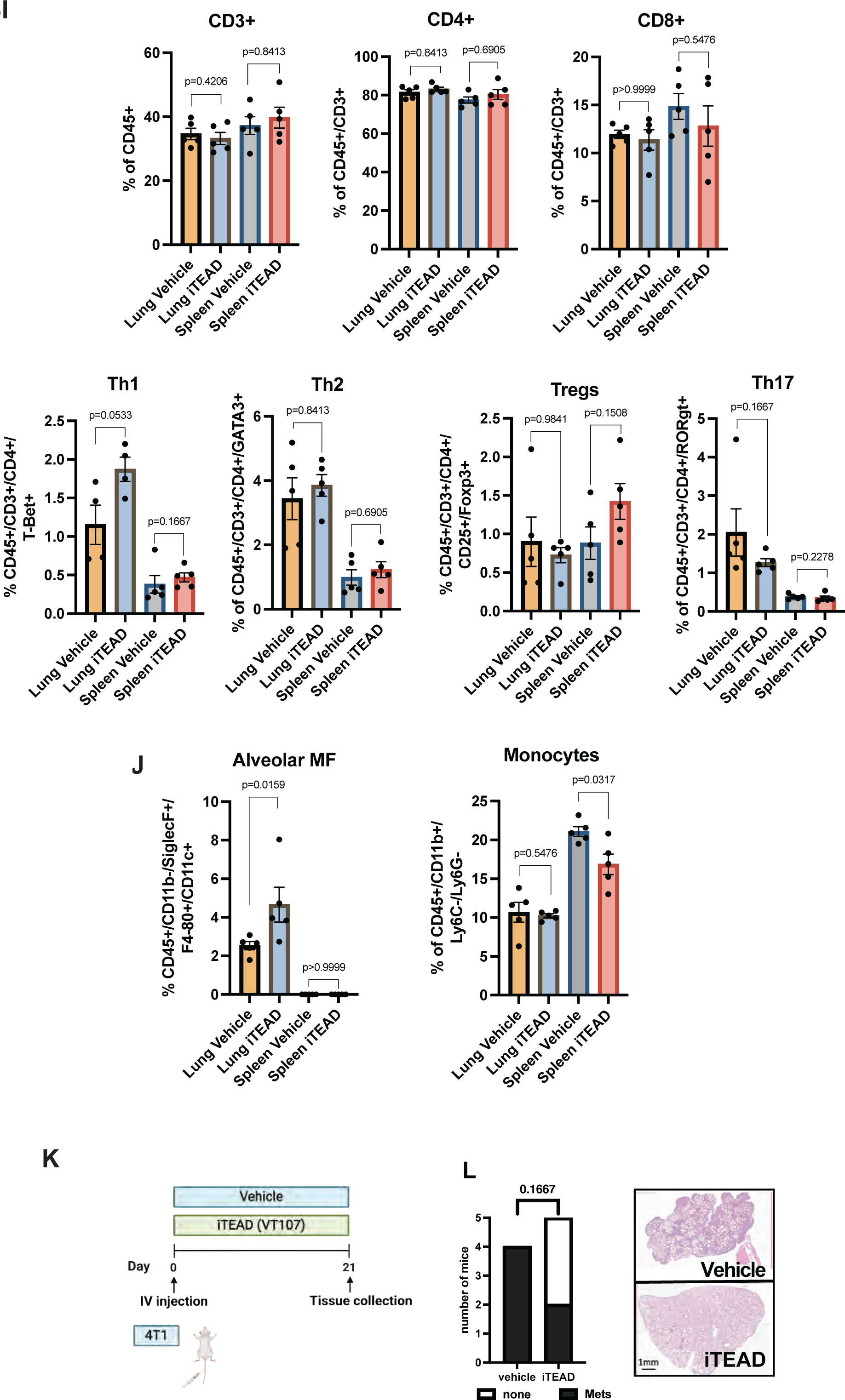
A) The body weight of C57/Bl6 mice bearing E0771 tumors with VT107 or vehicle treatment. B) RTCA of E0771 cells measured by electric impedance sensing of attached cells (cell index) over time with increasing doses of VT107. (Mean±SD, student t-test). C) The rate of tumor recurrence between treatment groups 2 weeks after E0771 tumor resection at week 4. (Student t-test). D) Volcano plots depicting change in tumor and lung gene expression of VT107-treated versus the control group. Red and blue circles indicate regulated genes with −Log p-value >1.3 and |log_2_FC|=2. A full list of changed genes is in supplementary table 1. E) Flow cytometry analysis of the alveolar macrophage population in the lung of mice that were treated with VT107 versus vehicle. (Mean±SD, student t-test). F) Flow cytometry analysis of the monocyte population in the lung and spleen of mice that were treated with VT107 versus vehicle. (Mean±SD, student t-test). G) Quantification of IHC of CD206 positive cells, an M2-like marker, in the lungs of VT107-treated versus the control group. (n=5 fields/lung). Images are representative of the lungs in each treatment group. H) Quantification of IHC of mono/macrophage marker CD68 and T cell marker CD8 in the tumors of VT107-treated versus the control group. (n=5 fields/lung, student t-test). I) Flow cytometry analysis of the lymphoid cells and T cell subtypes in the lung of mice that were treated with VT107 versus vehicle from panel 4H. (Student t-test). J) Flow cytometry analysis of the alveolar macrophage and monocyte population in the lung of mice that were treated with VT107 versus vehicle. (Student t-test). K) Schematic of BALB/c mice that were injected intravenously (IV) with 4T1 cells and treated with either vehicle or 30mg/kg VT107 daily by oral gavage for 21 days. L) Quantification of the number of BALB/c mice that developed lung metastases was determined by the histopathological analysis of H&E lung sections in the VT107 treated versus control group. Representative H&E-stained lung sections showing metastases. (Fisher’s exact test).

**Figure S4:**
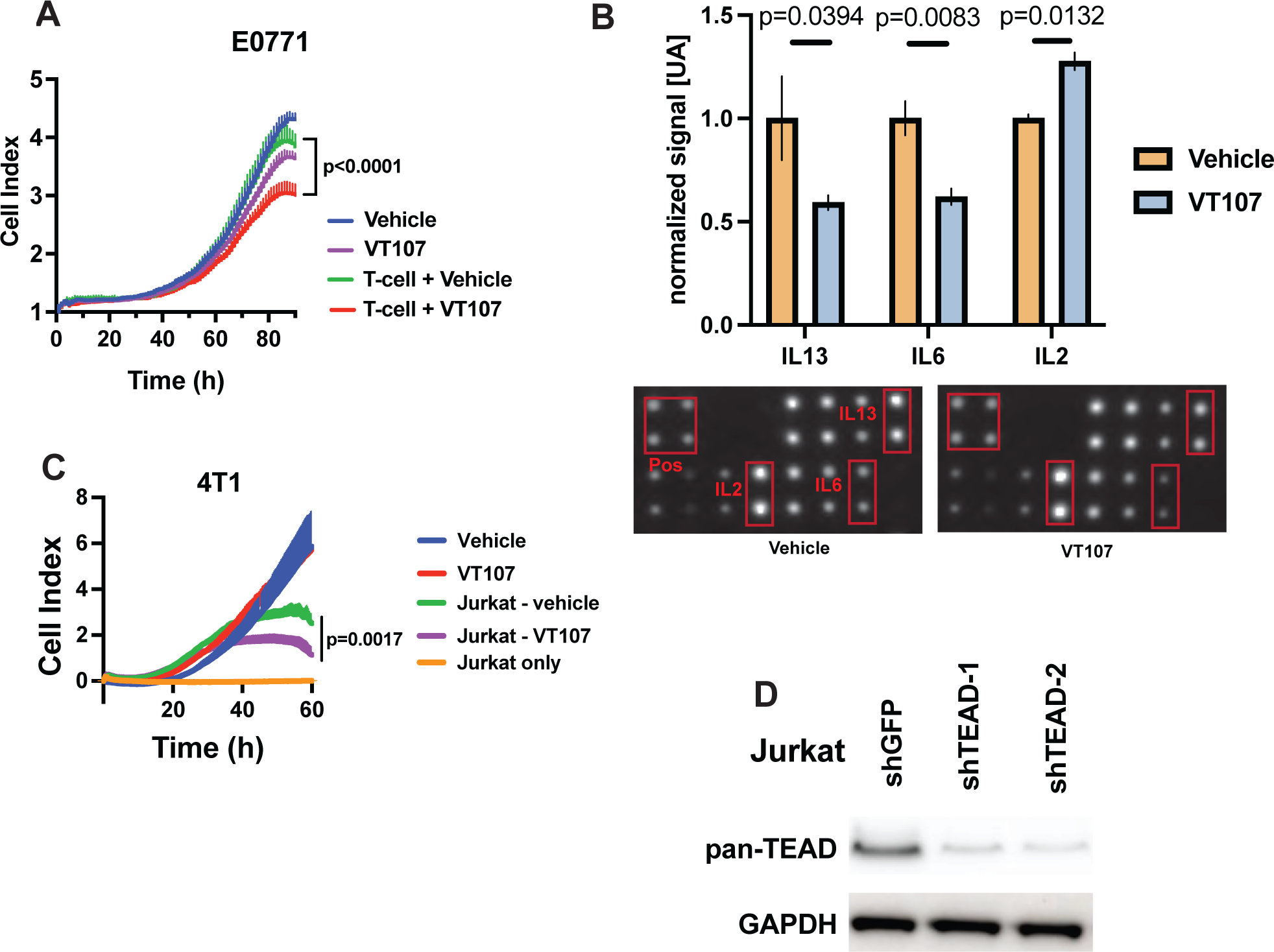
A) RTCA of E0771 cancer cells measured by electric impedance sensing of attached cells (cell index) over time. T cells treated with vehicle or 1μM VT107 for 24 hours were added simultaneously to the cancer cells to monitor their cytotoxicity. (Mean±SD, student t-test). B) Blot of cytokine array and quantified data showing changes in secreted factors in the conditioned media of T cells isolated from the spleen then treated with vehicle or 1μM VT107 for 72 hours. (Mean ± SEM, student t-test). C) RTCA of 4T1 cells was measured by electric impedance sensing of attached cells (cell index) over time. Jurkat cells treated with vehicle or 1μM VT107 for 24 hours then added to the cancer cells to monitor their cytotoxicity. (Mean±SD, student t-test). D) Western blot of pan-TEAD protein levels in Jurkat cells infected with either control shRNA or two distinct shTEAD lentiviruses. GAPDH protein levels were used as a loading control.

**Figure S5:**
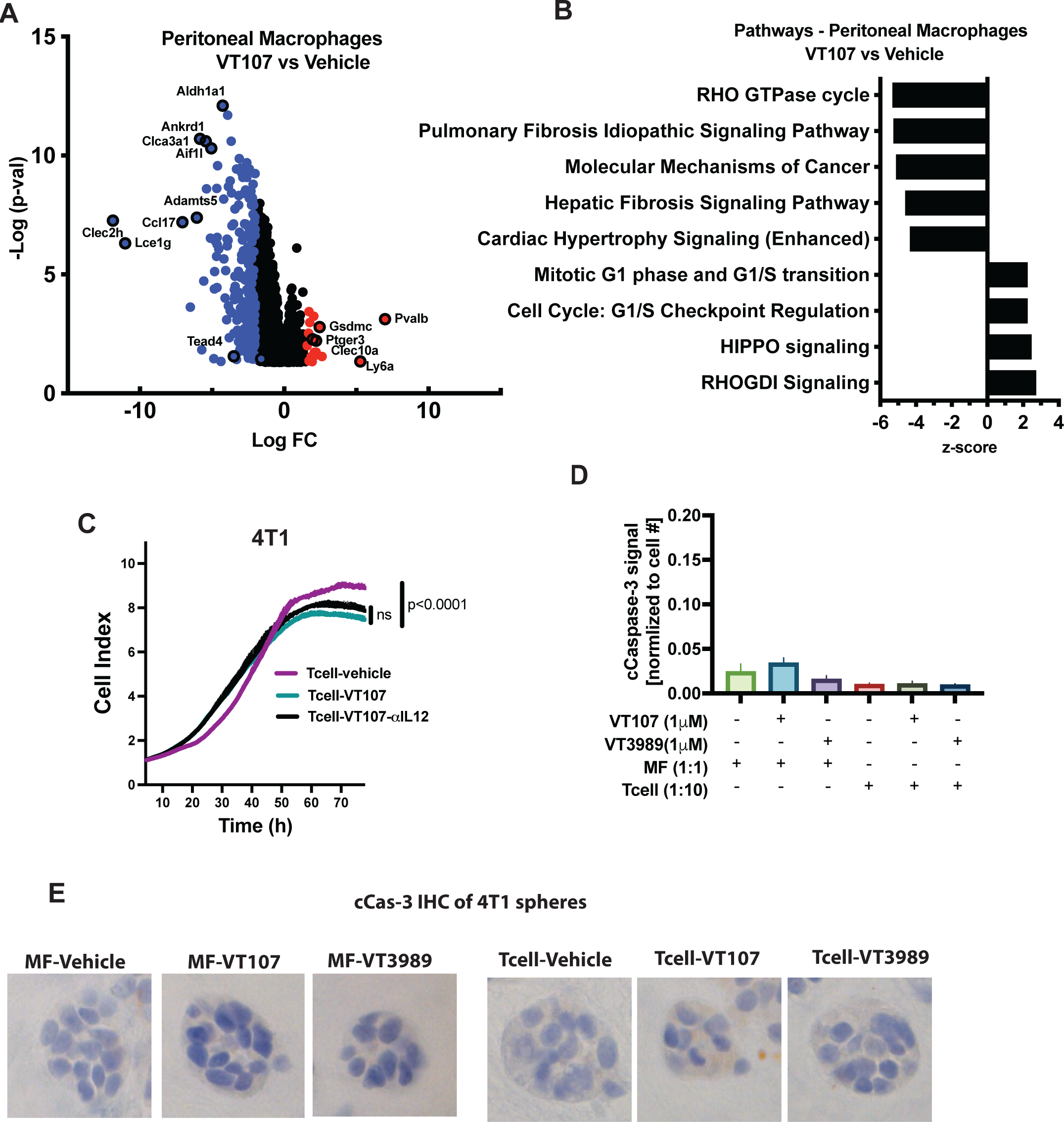
A) Volcano plot of the altered gene expression in peritoneal macrophages that were treated in vitro with vehicle or 1μM VT107 for 72 hours. Red and blue circles indicate regulated genes with −Log p-value >1.3 and |log_2_FC|=2. A full list of changed genes in supplementary table 2. B) IPA showing altered signaling pathways in peritoneal macrophages that were treated with vehicle or 1μM VT107 for 24 hours. −Log p-value >1.3 and |log_2_FC|=2. A full list of changed genes in supplementary table 2. C) RTCA of 4T1 cancer cells measured by electric impedance sensing of attached cells (cell index) over time. T cells treated with 1μM VT107 for 24hours were added simultaneously with the cancer cells to monitor cytotoxicity, in the presence or absence of 0.1 mg/ml IL12 antibody. (Mean±SD, student t-test). D) Quantification of the cCas-3 IHC signal 24 hours after adding the immune cells to the 4T1 spheres. n=3-10 spheres per condition. (Mean±SD). E) Representative images of IHC of cCas-3 on 4T1 spheres embedded in Matrigel/ collagen I mix. T cell and peritoneal macrophages were added to the spheres with the indicated treatments (vehicle, 1μM VT107,VT3989)

Supplementary Table 1: List of regulated genes and signaling pathways in mice (NOD/SCID, C57BL/6)

Supplementary Table 2: List of regulated genes, signaling pathways and upstream regulators in cell lines (THP1, Jurkat, peritoneal MF)

